# Alevin-fry unlocks rapid, accurate, and memory-frugal quantification of single-cell RNA-seq data

**DOI:** 10.1101/2021.06.29.450377

**Authors:** Dongze He, Mohsen Zakeri, Hirak Sarkar, Charlotte Soneson, Avi Srivastava, Rob Patro

## Abstract

The rapid growth of high-throughput single-cell and single-nucleus RNA sequencing technologies has produced a wealth of data over the past few years. The available technologies continue to evolve and experiments continue to increase in both number and scale. The size, volume, and distinctive characteristics of these data necessitate the development of new software and associated computational methods to accurately and efficiently quantify single-cell and single-nucleus RNA-seq data into count matrices that constitute the input to downstream analyses.

We introduce the `alevin-fry` framework for quantifying single-cell and single-nucleus RNA-seq data. Despite being faster and more memory frugal than other accurate and scalable quantification approaches, `alevin-fry` does not suffer from the false positive expression or memory scalability issues that are exhibited by other *lightweight* tools. We demonstrate how `alevin-fry` can be effectively used to quantify single-cell and single-nucleus RNA-seq data, and also how the spliced and unspliced molecule quantification required as input for RNA velocity analyses can be seamlessly extracted from the same pre-processed data used to generate regular gene expression count matrices.

## 1. Introduction

Both the number and scale of single-cell RNA-seq (scRNA-seq) experiments have been growing rapidly in recent years (1). The data generated by various single-cell sequencing technologies have distinct characteristics preventing them from being processed by the otherwise mature and widely used tools developed for gene-level or transcript-level quantification from bulk RNA-seq data (e.g. (2–4)). While `Cell Ranger` exists as a commercial solution for preprocessing data generated using the popular 10x Genomics technologies platform, it is both computationally and memory intensive, and since version 3 has been developed as a closed-source product, limiting the transparency of the methods it implements. Further, it does not have built-in support for technologies beyond those developed by 10x Genomics. Therefore, to address the computational challenges that arise in the processing of high-throughput single-cell RNA-seq data, numerous new approaches for efficient preprocessing have been developed.

Srivastava et al. (5) introduced `alevin`, which focused on improving the computational efficiency of tagged-end single-cell RNA-seq quantification and also introduced a novel approach for resolving gene-multimapping UMIs. Likewise, the `raindrop` tool (6) pairs a custom lightweight mapping approach with a reduced index to count UMIs mapping to genes, providing a fast counting approach. Melsted et al. (7) introduced the `kallisto|bustools` pipeline for processing single-cell RNA-seq data; the approach focuses on modularity and speed, using pseudoalignment (3) to the transcriptome to produce intermediate BUS files (8) that are subsequently manipulated using `bustools` commands.

Most recently, Kaminow et al. (9) introduced `STARsolo`. This preprocessing method is built directly atop the `STAR` aligner on which `Cell Ranger` also relies. `STARsolo` focuses on being a fast and easy-to-use solution for processing single-cell and single-nucleus RNA-seq (snRNA-seq) data. While being much faster and more memory frugal than `Cell Ranger`, it can be tuned to mimic `Cell Ranger` almost exactly. However, by virtue of performing spliced alignment to the genome, `STARsolo` is more memory and time-intensive than pseudoalignment to the transcriptome (at least for scRNA-seq data).

In this work, we present `alevin-fry`, a configurable framework for the processing of scRNA-seq and snRNA-seq data. `Alevin-fry` has been designed as the successor to `alevin`. It subsumes the core features of `alevin`, while also providing important new capabilities and considerably improving the performance profile, and we anticipate that new method development and feature additions will take place primarily within the `alevin-fry` codebase. `Alevin-fry` can preprocess scRNA-seq data much more quickly than the next-fastest method, `kallisto|bustools`, while also virtually eliminating the large numbers of spuriously expressed genes predicted under pseudoalignment-to-transcriptome approaches (9). Simultaneously, `alevin-fry` exhibits similar accuracy to `STARsolo` while processing data appreciably faster and requiring less memory. On snRNA-seq data, where intronic sequences are often included for quantification, `alevin-fry` and `STARsolo` are both faster and use less memory than `kallisto|bustools`. In fact, `alevin-fry` can process snRNA-seq data with the same speed and memory efficiency with which it processes scRNA-seq data, substantially outperforming both `STARsolo` and `kallisto|bustools`. `Alevin-fry` is an accurate, computationally efficient, and easy-to-use tool that presents a unified framework for preprocessing single-nucleus and single-cell RNA-seq data for gene expression or RNA velocity analysis, making it an appealing choice for processing the diverse and growing array of single-cell RNA-sequencing experiments being performed.

## 2. Methods

`Alevin-fry` is a configurable framework for the processing of single-cell and single-nucleus data. It makes use of `salmon` (4) for basic barcode and UMI parsing and the mapping of the reads to the constructed reference index. The output of `salmon`, when configured to produce output for `alevin-fry`, is a RAD (Reduced Alignment Data) format file, which is a chunk-based binary file optimized for machine parsing, that encodes the relevant information necessary for subsequent (post-mapping) processing of the data (Section S1). `Alevin-fry` consumes the `salmon` output directory — containing the RAD file and other relevant meta information about the sample — and processes the data in a number of steps. The main processing steps correspond to permit-list generation, RAD file collation, and finally, quantification of the collated RAD file. We describe the options provided by `alevin-fry` and further details of these specific steps below.

### 2.1. Constructing a reference index

The `alevin-fry` workflow quantifies single-cell data based on a reference index created by `salmon`. Here, we will discuss two types of reference sequences that can be used to construct such an index, and describe the relative advantages and disadvantages of these options. Regardless of the reference over which one decides to build an index, `salmon` makes use of the pufferfish (10) index, and a dense or sparse index variant can be constructed.

First, at least for the processing of single-cell (not single-nucleus) data, one might consider building a reference index over the spliced transcriptome. The main benefits of this approach are that it is simple, and the resulting index tends to be very small. For example, when using the spliced transcriptome extracted from the latest 10x Genomics version of GRCh38, the (dense) reference index is only ∼ 700MB, and the entire mapping and quantification procedure can be performed in ∼ 3GB of RAM.

However, while the frugal resource use of an index restricted to only the spliced transcriptome is appealing, it comes with potential drawbacks. The most significant drawback, perhaps, is that it results in substantial false positive rates (i.e. spuriously detected genes) (9). One likely mechanism is that in typical single-cell experiments, some fraction of reads (in cases, up to ∼ 25%) derive from intronic or intergenic sequences rather than from spliced transcripts (9). When these true sequences of origin are absent from the index, reads deriving from them may sometimes be spuriously assigned to a spliced transcript that shares some local sequence similarity with the true locus of origin. The degree to which such spurious assignment occurs also depends on the specifics of the algorithm used for mapping; for example, the problem is most pronounced when using pseudoalignment (3), followed by pseudoalignment with structural constraints, and is somewhat (but not fully) mitigated when using selective-alignment (9).

One alternative is to map to the genome directly, as is done by `Cell Ranger` and `STARsolo`. This allows consideration of all genomic loci when determining the appropriate mapping location for a read, and results in the elimination of the false positives that are induced by forcing reads to map only against the annotated transcriptome. While building such an index is quite comprehensive, the associated costs are that the index is inevitably larger, and the common alignment approaches for single-cell data (both `Cell Ranger` and `STARsolo` are based on STAR (11) as their underlying aligner) require considerably more RAM during alignment. Further, these approaches require solving the spliced (rather than contiguous) alignment problem; while good solutions (like STAR and HISAT2 (12)) exist, this problem is more computationally intensive and lightweight approaches like quasi-mapping (13) and pseudoalignment (3) have not yet been adapted to the problem of spliced mapping.

We propose here an alternative middle-ground, which is to align against a reference that indexes both the spliced transcriptome and the set of (collapsed) intron sequences that are likely to generate reads in a typical single-cell (or single-nucleus) experiment. We employ a reference preparation algorithm to produce what we refer to as a *splici* (spliced + intronic) reference, representing a slight modification of references previously used for RNA velocity preprocessing (7, 14). Further details explaining how this reference is constructed are provided in Section S2. Unlike the spliced transcriptome alone, this index contains the intronic sequences that are likely to give rise to a non-trivial fraction of reads in a single-cell experiment, and including these sequences allows one to properly resolve read origin and avoid the spurious mapping associated with mapping against the spliced transcriptome alone, similar to what is accomplished by decoy sequences in bulk RNA-seq quantification (15) (though different in execution, as the quantification method itself, and not just the mapping algorithm, is aware of these sequences). On the other hand, by indexing the spliced transcriptome and introns (with flanking sequence) directly, this reference does not require spliced alignment and is therefore amenable to both fast contiguous alignment algorithms like selective-alignment (15) as well as lightweight approaches like pseudoalignment (3). Throughout this manuscript, we set the flank length as the read length minus 5, though the quantification results appear very robust to the specific flank length chosen Section S3. While the size of this reference is considerably larger than the spliced transcriptome alone, it is still smaller than the genome. For the index used by `alevin-fry`, a dense index for a recent human reference constructed in such a manner requires ∼ 10GB of RAM for mapping, while the sparse index requires only ∼ 6.5GB. We demonstrate below how mapping against this index addresses the shortcomings of mapping against just the spliced transcriptome, while retaining modest memory requirements.

### 2.2. Fragment mapping

As with constructing a reference against which to map reads, multiple choices can be made as to exactly how fragments should be mapped to the reference. In `alevin-fry` there are two main options available; selective-alignment (15), and pseudoalignment (3) with structural constraints.

Broadly, selective-alignment is more accurate but more computationally intenstive. Fragments are mapped against the index using maximal exact matches between reads and indexed unitigs (uniMEMS) as seeds, which are then chained to determine a putative mapping score. Low-scoring putative mappings are discarded, and high-scoring mappings are validated using alignment scoring via dynamic programming, based on the banded, parallel implementation of minimap2 (16). All best-scoring alignments that are above a userdefined threshold are reported as valid alignments for the fragment. The explicit alignment scoring avoids the reporting of mappings where the locus having the best set of seed matches is not the locus having the best alignment. Likewise the discarding of alignments below the user-defined threshold ensures that fragments arising from some other origin that have no high-quality alignment in the indexed reference will not be reported and processed as valid mappings.

On the other hand, pseudoalignment with structural constraints, exposed via the `--sketch` flag, is very fast, but it does not validate mapping locations via alignment scoring. This approach first uses a custom implementation of pseudoalignment (3) to determine which k-mers from the fragment match different targets. Subsequently, the implied mappings are subjected to filtering by structural constraints requiring that the matches supporting the pseudoalignment are in a consistent orientation, are co-linear with respect to the read and the reference, and that the stretch (maximum distance between any pair of k-mers comprising the mapping) is not too large. While using a *splici* index largely eliminates the problem of false positive expression that has previously been reported when using pseudoalignment-to-transcriptome approaches (9), enabling accurate quantification using this rapid approach, there are still some false positive mappings that can only be properly eliminated with alignment scoring (i.e. using selective-alignment).

### 2.3. Permit-list generation

After the reads have been mapped to the target index, either using selective-alignment or pseudoalignment with structural constraints, the resulting RAD file is inspected to determine the set of cellular barcodes (CBs) that should be used for quantification. In single-cell sequencing experiments, cell capture rates are imperfect, and thus some fraction of barcodes may correspond to droplets that failed to properly capture a cell (17). In this case, the fragments associated with these barcodes usually exhibit many fewer distinct Unique Molecular Identifiers (UMIs) mapped to target sequences in the index than barcodes corresponding to properly captured cells. Likewise, errors that occur during PCR amplification and sequencing can “corrupt” the sequence of a cellular barcode, so that the barcode observed in the sequenced fragment is different from that which was originally attached to the underlying molecules prior to sequencing.

`Alevin-fry`’s `generate-permit-list` command works to determine the set of CBs that will eventually be quantified, as well as to perform correction of likely-corrupted barcodes to the “true” barcode from which they derived. It exposes a number of different strategies to determine the set of CBs that should be quantified. The supported strategies are `--force-cells`, `--expect-cells`, `--knee-distance`, `--unfiltered-pl` and `--valid-bc`. Here we briefly describe the `--knee-distance` and `--unfiltered-pl` strategies, since they are likely to be the most commonly employed by users of `alevin-fry`. A description of all available methods is provided in the `alevin-fry` documentation (https://alevin-fry.readthedocs.io/en/latest/generate_permit_list.html). This step is also used to apply orientation filtering to the mapped records. So, for example, in protocols where all fragments are expected to map to the reference in the forward orientation, fragments (and their associated barcodes) are only considered valid if at least one forward strand mapping exists.

#### Knee distance permit-list generation

The knee distance filtering implemented in `alevin-fry` is an implementation of the (updated) strategy that is provided in the `UMI-tools` (18) software. It is an iterative knee finding strategy that attempts to automatically determine the number of barcodes corresponding to high-quality cells by examining the frequency histogram of observed barcodes. Briefly, this method first counts the number of reads associated with each barcode, and then sorts the barcodes in descending order by their associated read count. It then constructs the cumulative distribution function from this sorted list of frequencies. Finally, it applies an iterative algorithm to attempt to determine the optimal number of barcodes to include by looking for a “knee” or “elbow” in the CDF graph. The algorithm considers each barcode in the CDF where its x-coordinate is equal to this barcode’s rank divided by the total number of barcodes (i.e. its normalized rank) and the y-coordinate is equal to the (normalized) cumulative frequency achieved at this barcode. It then computes the distance of this barcode from the line x = y (defined by the start and end of the CDF). The initial knee is predicted as the point that has the maximum distance from the x = y line. The algorithm is iterative, because experiments with many low-quality barcodes may predict too many valid barcodes using this method. Thus, the algorithm is run repeatedly, each time considering a prefix of the CDF from index 0 through the previous knee’s index times 5. Once two subsequent iterations of the algorithm return the same knee point, the algorithm terminates. Once the set of “permitted” barcodes has been determined by this method, the reads that have barcodes not within this set are corrected against it by checking if they are within one edit of some barcode in the list; if so, they are attributed to that barcode.

#### Correcting to an unfiltered permit-list

Some technologies, like 10x Chromium, provide a set of specific known and experiment-independent barcodes that will be a superset of the barcodes that should be observed in any given sample. This list of “possible” barcodes can be treated as a set of barcodes against which the observed barcodes can be corrected. The `--unfiltered-pl` option accepts as an argument a list of possible barcodes for the sample. When using this argument, the user may also pass the `--min-reads` argument to determine the minimum frequency with which a barcode must be seen in order to be retained. The algorithm used in this mode passes over the input records (mapped reads) and counts how many times each of the barcodes in the unfiltered permit list occurs exactly. Any barcode occurring ≥ min-reads times will be considered as a present cell. Subsequently, all barcodes that did not match a present cell will be searched (at an edit distance of up to 1) against the barcodes determined to correspond to present cells. If an initially non-matching barcode has a unique neighbor among the barcodes for present cells, it will be corrected to that barcode, but if it has no 1-edit neighbor, or if it has two or more 1-edit neighbors among that list (i.e. its correction would be ambiguous), then the record is discarded. Of course, unfiltered count matrices constructed in this manner will contain many barcodes not corresponding to properly captured cells, and should be subjected to subsequent filtering prior to analysis.

In all cases, the result of the `generate-permit-list` step of `alevin-fry` is the creation of a “correction map” that specifies which barcodes are to be quantified, and how barcodes are to be corrected against this quantified set, as well as a census of the number of observed and valid fragments corresponding to each *corrected* barcode. The census information is used in the subsequent collation step to enable an efficient partitioning strategy for collating the records by corrected barcodes.

### 2.4. Collation of RAD files

Once the permit-list and correction map have been generated, the initial RAD file must be *collated* by the corrected cellular barcodes; this is done using `alevin-fry`’s `collate` command. In this phase, all fragments to be quantified are grouped together such that those sharing the same barcodes appear contiguously in the file. This step of processing serves an analogous purpose as the sorting of BUS (8) files as done in `kallisto|bustools` (7). However, there are a few technical differences that relate to the way in which `alevin-fry` processes the collated RAD files and to the way in which RAD files are structured differently from BUS files.

First, the `collate` command, unlike sorting, does not induce a total order on the resulting records. Specifically, while records that pass filtering and have the same corrected barcode are guaranteed to occur contiguously within the resulting collated file, there is no specific or meaningful order between the segments of the file corresponding to individual corrected barcodes. Further, within the set of records corresponding to a corrected barcode, there is no ordering or collation among the UMIs. This is due, in part, to the fact that all records sharing the same corrected barcode will be present in memory at the same time during the quantification phase, as well as the fact that certain UMI resolution strategies apply edit-distance collapsing for which UMI collation is insufficient. This means that, in the general case, collation can potentially be implemented in a more computationally efficient manner than sorting, though in practice, multithreaded sorting of fixed sized records is very efficient.

Eliminating the requirement of having sorted records also means that the collated records corresponding to each barcode can appear at whichever location in the collated output file that is desired. This admits extra flexibility in how collation is performed. Specifically, corrected barcodes are assigned to roughly appear in order of descending frequency in the output collated RAD file. This means that the largest (and potentially slowest to process) cells will appear near the beginning of the collated RAD file. Since quantification itself is multithreaded, this allows more efficient pipelining of quantification among multiple threads. Since threading granularity happens at the level of individual cells (i.e. the records for the same cell will never be quantified by multiple threads at the same time), placing the largest cells early in the quantification phase means that one is unlikely to encounter a situation where large and complex cells are encountered late in processing and many threads remain work starved while processing for the large cell completes.

The collation strategy implemented in `alevin-fry` is a two-pass approach. First, each corrected barcode is assigned a bucket index; the input RAD file is parsed (in parallel by many worker threads), and each record is written to the bucket it is assigned based on its corrected barcode. This ensures that all records sharing the same corrected barcode are routed to the same bucket. Further, the bucket sizes are limited by a user-defined maximum record count to ensure that individual buckets can be fully loaded into memory while retaining an overall small memory profile. In a second pass, each bucket is read into memory and its records are locally collated. This is done by constructing an in-memory hash map mapping each corrected barcode in this bucket to the vector of records sharing this barcode. Subsequently, each locally collated chunk is appended to the output collated RAD file (and optionally compressed if the user passes the `--compress` flag). In the resulting RAD file, the number of chunks is equal to the number of cells to be quantified (i.e. the number of corrected barcodes), and all of the records sharing the same corrected barcode appear consecutively in the file.

### 2.5. Quantification

With the collated RAD file prepared, `alevin-fry` is able to quantify the count for each gene within each cell separately and in parallel via the `quant` command. As with the mapping and permit-list generation phase, a number of different UMI resolution strategies are implemented in `alevin-fry`. Here, we briefly describe those strategies — `cr-like` and `cr-like-em` — that currently support the unspliced, spliced and ambiguous (USA) quantification mode that is used throughout this manuscript. As opposed to splice unaware quantifciation (which `alevin-fry` also supports), the USA quantification mode produces a count for each splicing status of each gene within each quantified cell. Additional resolution strategies are described in Section S4.

The quantification for each cell is carried out independently and in parallel, so we explain the procedure, without loss of generality, for the records corresponding to an individual cell. First, read records are collated (in memory) by their corresponding UMI. For each UMI, the set of transcripts to which the read maps is projected onto the corresponding set of genes. This process is aided by the use of a three element transcript to gene map. Each entry in the map contains the name of an individual target sequence from the splici index, the corresponding gene to which this target belongs, and a splicing status, recorded as ‘S’ if the target derives from a spliced transcript and ‘U’ if it derives from intronic sequence. Each gene is assigned a pair of globally unique identifiers, one corresponding to all “spliced” variants of the gene and the other to the “unspliced” (intronic) sequences for the gene. The gene-level identifiers corresponding to a given record are sorted and deduplicated. All records corresponding the current UMI are iterated in the same fashion, and a count is kept of how many times the UMI is associated with a read that maps to each gene identifier (with “spliced” and “unspliced” identifiers treated as distinct).

After all occurrences of the UMI are observed, the UMI is assigned to the gene with the largest frequency. If there is no unique gene with the highest frequency of occurrence, then the UMI is discarded if the `cr-like` resolution strategy is being used. On the other hand, if the `cr-like-em` resolution strategy is being used, a gene-level equivalence class is formed from all gene identifiers having the highest frequency of mapping for this UMI. Each identifier in the label of the equivalence class comprises a gene and a splicing status. The status is ‘U’ if only the unspliced identifer of this gene is among the most frequent mapping targets for this UMI, it is ‘S’ if only the spliced identifier is among the most frequent, and if both the unspliced and spliced identifiers of this gene are among the most frequent mapping targets for this UMI, then the status is ‘A’ (ambiguous). The UMI is attributed to this equivalence class, and an expectation maximization (EM) algorithm, like that employed in Srivastava et al. (5), is subsequently used to probabilistically allocate counts to specific gene and splicing status pairs in the resulting count vector for this cell.

Under both of these resolution strategies, the resulting count matrix contains a count not just for each gene within each cell, but the count is further distributed over each gene’s splicing status (confidently assigned to spliced molecules from the gene, confidently assigned to unspliced molecules from the gene, or ambiguous in splicing status). Depending upon the type of data analysis being performed, this count matrix can then be used to directly extract the counts of interest. For example, if performing a “standard” single-cell gene expression analysis, one can extract the spliced and ambiguous counts for each gene in each cell and sum them together to produce the equivalent of a standard count matrix. If performing quantification on a single-nucleus RNA-seq sample, the counts from all splicing categories can be summed to produce the total UMI count attributed to each gene (regardless of splicing status). For an RNA velocity analysis, the spliced and unspliced counts can be separated into distinct matrices and provided to a downstream RNA velocity computation tool (19, 20).

These resolution strategies thus provide a convenient solution for quantification of gene expression in a variety of different single-cell settings. The same processing approach can be used for the quantification of gene expression in single-cell experiments, or in single-nucleus experiments, or even to provide the input for RNA velocity analysis. At the same time, explicitly accounting for the unexpected origin of reads (e.g., from intronic sequence in single-cell experiments) can also virtually eliminate the spurious detection of genes exhibited by methods that restrict mapping or alignment to only the spliced transcriptome. This is possible as these resolution strategies implemented by `alevin-fry` are designed to infer both the gene and splicing status of the underlying fragments, but leave the determination of how to combine or aggregate UMIs arising from different splicing statuses to downstream analysis.

Finally, a number of additional and even more sophisticated resolution methods (namely `parsimony` and `parsimony-em`) are present in `alevin-fry` but not yet exposed under USA mode. These implement variants on the original UMI resolution algorithm introduced by Srivastava et al. (5) that applies a parsimony condition to approximately determine the minimal set of transcripts that could give rise to the observed set of UMIs. These alternative methods are further described in Section S4. We are currently working on adapting these algorithms so that they can also be meaningfully applied in `alevin-fry`’s USA mode.

### 2.6. RNA velocity

With the development of single-cell RNA-seq technologies, RNA velocity analysis has become increasingly popular. `Velocyto` (19) defines single-cell RNA velocity as the time derivative of the gene expression state, which is determined by the ratio of the spliced and unspliced mRNA transcript molecule counts of each individual gene. By modeling transcriptional dynamics, RNA velocity can reveal cellular differentiation dynamics and developmental lineages present in a given single-cell experiment. `scVelo` (20) further enhances RNA velocity computation by eliminating the steady-state assumption made by `Velocyto`, and applying an expectation maximization method to solve the differentiation dynamics according to a series of master equations. The accurate and robust estimation of RNA velocity remains an active and exciting area of research.

To explore preprocessing for RNA velocity analysis, we make use of a mouse pancreatic endocrinogenesis dataset introduced by Bastidas-Ponce et al. (21) and used as an example dataset in the `scVelo`. This experiment is obtained with the 10x Genomics Chromium Single Cell 3’ Reagent Kit v2 and the read length is 151 nt. To utilize the cell state annotation information provided in `scVelo` example dataset, only the 3,696 cells that are included in the `scVelo` example dataset are included in our analysis. The quantified cells are all from stage E15.5. The processing was performed on raw FASTQ files retrieved from the Gene Expression Omnibus (GEO), accession number GSM3852755.

Following the preprocessing steps adopted by `scVelo`, we downloaded the pre-built mouse *mm10 v2*.*1*.*0* reference sequences and GTF files from 10X Genomics. To obtain the appropriate input for RNA velocity analysis with `alevin-fry` we make use of USA mode quantification, `kallisto|bustools` was run via the `kb_python` tool with the `--workflow lamanno` option, which results in the generation of two separate output matrices corresponding to the spliced and the unspliced counts, and `STARsolo` was run with the `--soloFeatures Gene Velocyto` option.

Depending on the RNA velocity method being used, ambiguous counts (which are output separately by `STARsolo` and `alevin-fry`) should either be provided explicitly, or allocated among the spliced and unspliced counts (or discarded entirely). We tested 7 different strategies to process the ambiguous counts, which are for each gene within each individual cell, (i) discarding the ambiguous counts, (ii) regarding the ambiguous count as spliced, (iii) regarding the ambiguous count as unspliced, (iv) evenly distributing the ambiguous count to spliced and unspliced, (v) dividing the ambiguous count by the ratio of confidently spliced count to the confidently unspliced count, (vi) dividing the ambiguous count by the ratio of not-unspliced (spliced + ambiguous) to unspliced, and (vii) dividing the ambiguous counts by the ratio of spliced to not-spliced(unspliced + ambiguous). We discuss the results of approach (ii) in Section 3.3, and provide all other results in Section S6.

`ScVelo` version 0.2.3 is used to analyze RNA velocity under a python 3.8.5 environment. Cells whose cell barcode is in the `scVelo` example dataset are kept for further analysis. The pre-defined cell type and UMAP representation of each cell are obtained from the `scVelo` example dataset. The count matrices generated by all three methods are processed as described in Bergen et al. (20). Specifically, the count matrices are median normalized, only the top 2, 000 variable genes are kept, the first- and second-order moments of the normalized spliced and unspliced counts of each gene are calculated, and the reaction rates and latent variables are recovered. RNA velocity is estimated using the dynamical mode, and the directional flow of the estimated velocity is visualized in the predefined UMAP (22) embedding.

### 2.7. Clustering analysis of snRNA-seq data

To evaluate the process of quantifying an snRNA-seq dataset using these preprocessing tools, we performed a cell type clustering analysis on the data from Marsh and Blelloch (23) using `Seurat` 4.0.1 (24) under an `R` 4.0.5 environment. When analyzing single-nucleus RNA-seq data, we sum the unspliced (U), spliced (S), and ambiguous(A) counts returned by `alevin-fry` to get the overall count of each gene within each cell. Likewise, `kallisto|bustools` is run via the `kb_python` tool with the `--workflow nucleus` option specified, and `STARsolo` is run with the `--soloFeatures GeneFull` option.

A snRNA-seq mouse placenta dataset (23) is selected as the example for demonstrating single-nucleus clustering analysis. The nuclei were captured with the Chromium Single Cell 3’ Reagent V3 Kit from 10X Genomics, and the read length is 150nt. In this analysis, only the samples from day E14.5 are included. These raw reads can be accessed from GEO under accession code GSM4609872.

To compare the results from different quantification tools fairly, we implemented the `emptyDrops CR` functionality of `STARsolo` in `R`, and then applied this function to filter empty droplets for the results of all tools under the same setting, which is the default setting in `STARsolo`. Only barcodes whose FDR is less than 0.01, mitochondrial counts are less than 0.25%, and that have 500 − 4,000 unique genes were kept for further clustering analysis.

After filtering empty droplets, the RNA counts of each nucleus were log normalized. Next, the top 2,000 variable genes were detected and their gene counts were scaled and used in the following steps. Then, PCA was performed with those variable genes, and a subset of significant PCs were selected using the `JackStraw` algorithm implemented in `Seurat`. Using those PCs, the t-SNE dimensionality reduction (25) was calculated, the nearest neighbor graph was constructed and clustering was performed. In order to assign cell type to each cluster, the R object provided by Marsh and Blelloch (23) was used as the reference to transfer the cell type annotation from the reference samples to the query object according to the anchor genes determined by the significant PCs.

The nuclei assigned as trophoblast were then selected to explore the trophoblast subclusters. Similar to the previous procedure, clusters were found using a subset of significant PCs determined by `JackStraw`, and cell types were learned from the trophoblast R objected provided in the supplementary file of Marsh and Blelloch (23).

## 3. Results

Here we demonstrate the performance and accuracy of `alevin-fry` in a variety of different use cases, and compare its computational resource usage as well as the quality of its results to those provided by the other recently introduced tools `STARsolo` and `kallisto|bustools`. We examine results on simulated data (Section 3.1), on a specific single-cell RNA-seq dataset where the effect of alignment pipelines has previously been explored (Section 3.2), in the context of preparing count matrices for an RNA velocity analysis (Section 3.3), for the processing of single-nucleus RNA-seq data sets (Section 3.4), and finally we explore the overall runtime and peak memory usage characteristics on this broad array of datasets (Section 3.5).

### 3.1. Simulated data

We first evaluated the different methods on data from a non-parametric simulation first introduced by Kaminow et al. (9). This simulation is seeded with the PBMC5k experiment (26). We use the simulated data in which reads derive from across the genome at realistic rates (i.e. from introns, spliced transcripts, and intergenic sequences), but without the simulated gene-level multimapping. While not tied to any parametric model, and therefore likely to produce realistic mapping statistics, it is important to recall the caveat that simulated data often fails to recapitulate at least some important aspects of experimental data (15). This implies that the performance on simulated data likely represents, in a sense, the upper bound of accuracy achievable by these methods on experimental data, and the degradation in performance of different methods may vary as the complexity of the data increase. Nonetheless, analyzing the accuracy of these tools under various metrics on this simulated data provides an important perspective as to the potential strengths and shortcomings of different methods in a situation in which the ground truth counts are known.

Table 1 displays the results for the different methods as evaluated under various metrics on the set (intersection) of the cells quantified by all methods. The definitions of these metrics are given in Section S7. While no method yields the best performance universally, there are some clear trends that can be observed. First, as was noted by Kaminow et al. (9), the methods that perform mapping (either pseudoalignment or pseudoalignment with structural constraints) directly to the spliced transcriptome alone perform worse than the other approaches — often considerably — under almost all metrics (the sole exception being the mean per-cell relative false negative rate). Specifically, these approaches exhibit a markedly reduced cell-level Spearman correlation with the truth, as well as largely inflated relative false positive expression (27-32%) and increased mean absolute relative deviations (MARD).

**Table 1:** The performance of the examined tools under various accuracy metrics on the simulated data. All metrics are measures on the subset of genes and cells defined by all tested methods, and are taken with respect to ground-truth abundances.

| method | mean Sp. corr. | MARD (drop NA) | MARD (NA=0) | mean rFP/cell | mean rFN/cell |
| --- | --- | --- | --- | --- | --- |
| STARsolo | 0.997 | 0.031 | 0.002 | 0.001 | 0.005 |
| kallisto\|bustools | 0.864 | 0.263 | 0.024 | 0.328 | 0.006 |
| alevin-fry (txome, sketch) | 0.883 | 0.226 | 0.020 | 0.273 | 0.006 |
| alevin-fry (splici, sketch) | 0.988 | 0.026 | 0.002 | 0.011 | 0.012 |
| alevin-fry (splici, sla) | 0.992 | 0.019 | 0.001 | 0.004 | 0.011 |

To explore these false positive expression estimates in slightly more detail, Figure 2 shows the frequency distribution of the number of cells in which each gene appears, where genes are sorted in descending order independently per method. Specifically, we observe that `STARsolo`, and both variants of `alevin-fry` that make use of the *splici* index follow a very similar frequency distribution, and that this is distinct from the frequency distribution followed by `kallisto|bustools` and `alevin-fry` when mapping only to the transcriptome. This suggests that, not only does mapping to the transcriptome alone result in hundreds of spuriously expressed genes per cell, but many of these genes themselves are expressed across hundreds of cells. Among the two evaluated approaches that map only to the spliced transcriptome, `alevin-fry` (in `sketch` mode) performs better than `kallisto|bustools`. On the other hand, the methods that map to expanded references, either the whole genome in the case of `STARsolo` or the splici reference in the case of `alevin-fry`, all generally perform well under the various metrics. `STARsolo` exhibits the highest cell-level Spearman correlation, as well as the smallest relative false positive and relative false negative rate, while `alevin-fry` exhibits the lowest MARD (both when run in `sketch` mode and when using selective-alignment).

**Figure 1:**
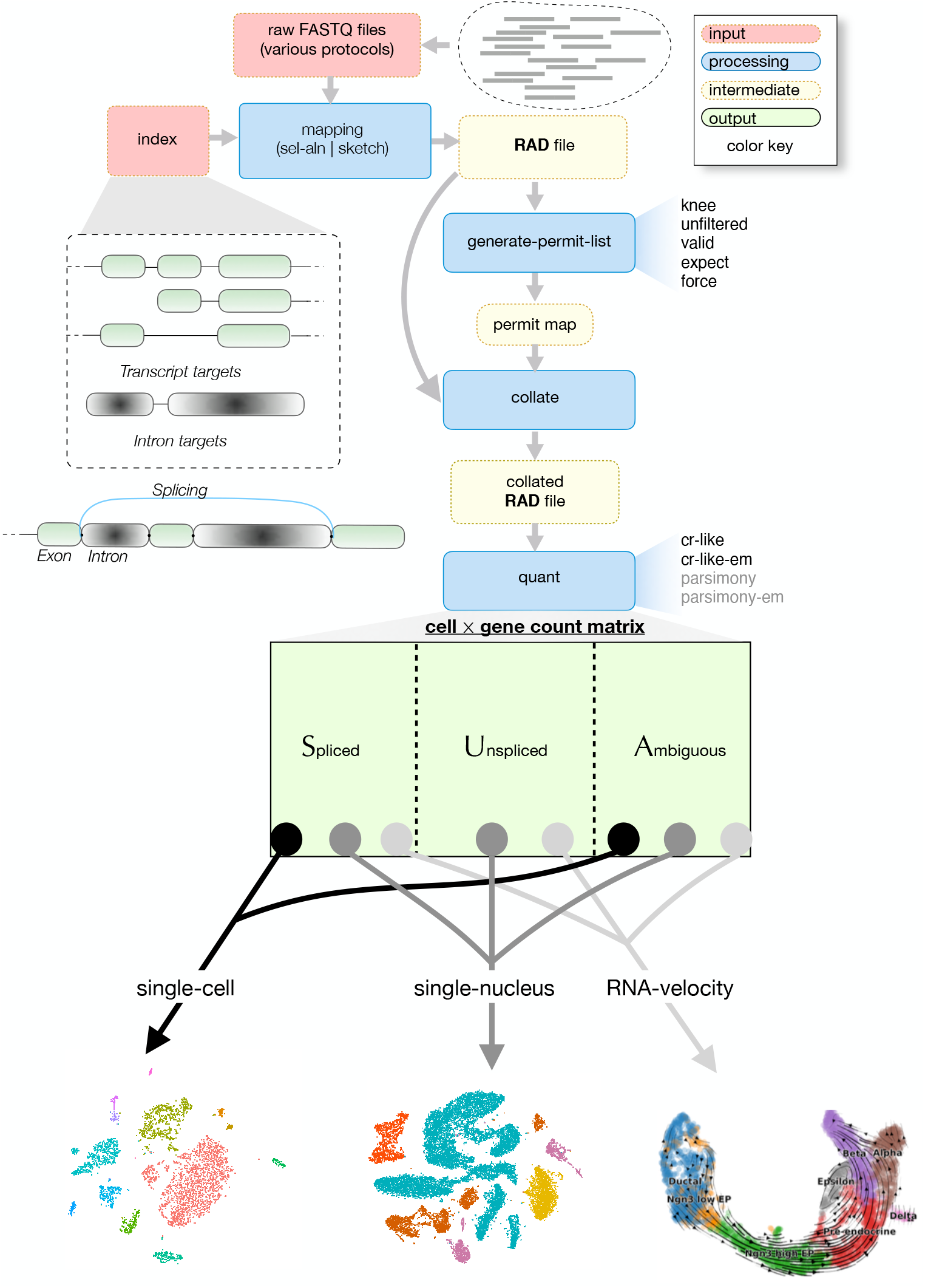
Overview of the `alevin-fry` pipeline (operating in unspliced, spliced, ambiguous quantification mode). The arrows highlight the flow of data through the pipeline, whose output is a matrix specifying the expected counts of each of the considered splicing states of each gene within each quantified cell.

**Figure 2:**
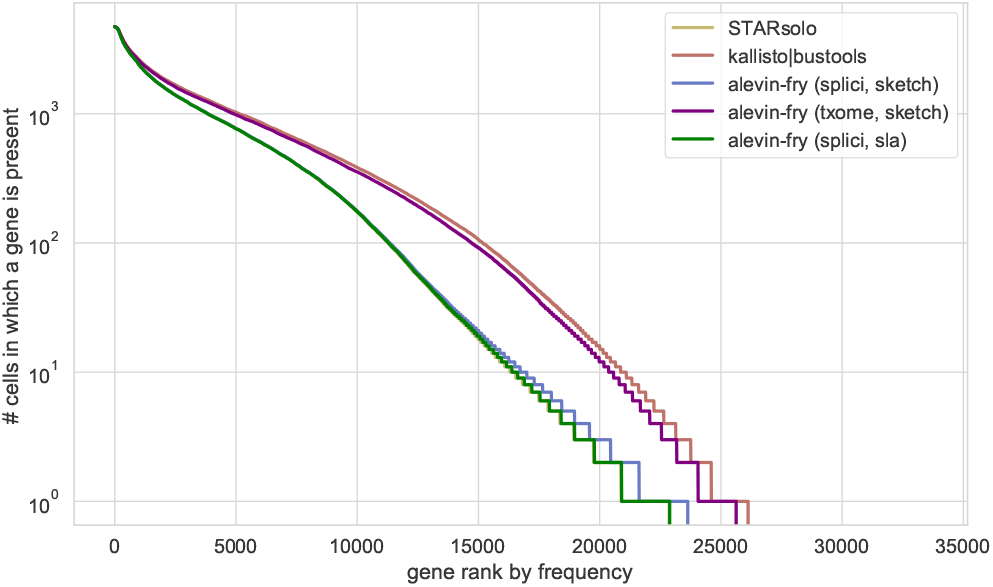
The frequency distribution of the presence of genes across all shared cells for `STARsolo`, `kallisto|bustools`, and `alevin-fry` (including multiple index types for `alevin-fry`).

Though there are differences under all metrics reported by the methods mapping to the expanded reference, the magnitude of these differences is generally small, and, in particular, is much smaller than the difference between any of these methods and those methods that map only to the spliced transcriptome. Moreover, we observe that, holding the other variables fixed (e.g. the reference and UMI resolution strategy), selective-alignment yields a small but consistent accuracy improvement over pseudoalignment with structural constraints. Presumably, this results largely from the ability of selective-alignment to discard fragments arising from outside of the spliced or unspliced transcriptome that would otherwise be spuriously assigned to some target. Nonetheless, we observe that pairing the expanded (*splici*) transcriptome with an appropriately UMI resolution strategy that is aware of both spliced and unspliced gene variants allows for the use of `sketch` mode (pseudoalignment with structural constraints) in a manner that corrects the high number of false positive expression predictions that are otherwise observed when mapping only to the spliced transcriptome.

### 3.2. Analysis of a *Danio rerio* pineal experiment

To explore the performance of `alevin-fry` in species other than human and mouse, and in an experimental sample where the alignment pipeline has previously been shown to have an impact on downstream analysis, we analyzed the *Danio rerio* pineal gland dataset introduced in Shainer et al. (27), and subsequently re-analyzed in (28).

Shainer and Stemmer (28) reported that for the *D. rerio* pineal dataset, `kallisto|bustools`’s result enables the `FindClusters` function of `Seurat` to recover an important biological feature that is not uncovered when examining the `Cell Ranger` output. Specifically, the resulting clustering uncovers two distinct *cone photoreceptor* (PhR) clusters, the cone PhRs expressing the *red cone opsin* (*red cone PhRs* or *red+* cells) and the cone PhRs expressing the *parietopsin* (*PT cone PhRs* or *PT+* cells). These two cell types, which are derived from a common PhR-restricted progenitor but that express distinct opsins, are two mutually exclusive neuronal fates of photoreceptors in this tissue (29). The *PT+* cells produce *parietopsin* (30), a green-sensitive photopigment that belongs to the non-visual opsins, whose expression is regulated by the repression of Notch activity and the stimulation of bone morphogenetic proteins (BMPs) activity. The *red+* cells generate *opn1lw1* (31), a long wavelength-sensitive (red) visual opsins expressed in *D. rerio* pineal tissue. On the other hand, when using `Cell Ranger`’s inferred counts, these two clusters are collapsed into a single *cone PhR* cluster that expresses the main marker genes for both the *red cone PhR* and *PT cone PhR* clusters. Interestingly, the strongest differentially expressed marker gene between the *red* and *PT cone PhR* clusters when analyzing the `kallisto|bustools` counts is *col14a1b*, a collagen gene.

To further investigate the differences demonstrated in (28) under a different set of pre-processing tools, we processed this data with `STARsolo`, `kallisto|bustools`, and `alevin-fry` using USA mode and the *splici* reference. Further, we ran `STARsolo` with a number of different UMI deduplication strategies — `1MM` (default), `1MMDir` and `exact` UMI deduplication strategies — the latter two being options not available in `Cell Ranger`. To normalize across the cell filtering method, we ran all tools to produce unfiltered quantifications, and subsequently filtered the resulting count matrices using the `DropletUtils` (17) R package, and we also evaluated the results of all methods on only the subset of cellular barcodes discovered by `alevin-fry`’s knee-finding method.

We first reproduced the analysis performed in (28), to observe the clustering that would result from the counts of `STARsolo` (using default UMI collapsing) and `alevin-fry`. In order to estimate the high-confidence cells from empty droplets, the `barcodeRank` function in `DropletUtils` was applied on the count matrices with a lower bound set to 500 to determine the `inflection` point on the UMI count of barcodes. Barcodes that have a UMI count below the `inflection` point were regarded as empty droplets and were filtered from the count matrices. The filtered count matrices were then used to create a `Seurat` object using the `CreateSeuratObject` function in `Seurat` with thresholds `min.cells = 3` and `min.feature = 200`. The resulting figures are provided in Figure S5.1 and Figure S5.2. We found, as described in (28), that `kallisto|bustools` exhibits distinct clusters for the *red* and *PT* cone PhR cells, and that the strongest differentially expressed gene (DEG) between these two clusters was *col14a1b*.

Likewise, we found that `STARsolo` using the default `1MM` strategy did *not* separate the relevant cells into distinct *red* and *PT cone PhR* clusters at a resolution parameter of either 0.9 or 1.2. This was expected given that `STARsolo`, under default parameter settings, is designed to closely mimic the approach and output of `Cell Ranger` (while being much more computationally efficient) (9). `Alevin-fry` (which was not considered in the original manuscript) does separate the relevant cells into two distinct clusters at both the 0.9 and 1.2 resolution parameters. However, we find that in the `alevin-fry` results, *col14a1b* is not one of the marker genes detected by `FindAllMarkers` function of `Seurat`. In fact, as with the `STARsolo` results, the *col14a1b* gene is not detected at any appreciable level among any of the cells after the prescribed filtering is applied. We investigated this difference and report our findings below and the clusterings and features plots in Section S5. However, we first wished to determine what differences might be causing the clustering of the `STARsolo` counts to fail to separate these cells into two distinct clusters when the clustering of both the `alevin-fry` and `kallisto|bustools` counts yields the distinct clusters.

One major difference is the `1MM` (default) method used for UMI deduplication, which attempts to mimic the algorithm used in `Cell Ranger`. To this end, we explored the effect of changing the default UMI resolution strategy applied by `STARsolo`, which is an option not exposed in `Cell Ranger`. We tested the `1MMDir` method that implements the directional algorithm described in UMI-tools (18), and the `exact` method that only deduplicates sequence identical UMIs. We discovered that under this filtering regime (28), the `STARsolo` counts when using both the `1MMDir` and `exact` deduplication methods yield distinct clusters for the *red* and *PT cone PhR* clusters at resolution parameters of both 0.9 and 1.2. Thus, in this data, the deduplication method seems to be an important factor for the signal separating these clusters to be picked up by `Seurat`’s clustering algorithm. It is worth noting that even when distinct clusters are not found, the t-SNE embedding computed from `STARsolo` counts places subsets of the cone PhRs in different positions in the embedding, and the subsets at these different positions express, disjointly, the marker genes for the *red+* and *PT+* cells; the clusters are just not separated by the clustering algorithm, as shown in Figure S5.2e and Figure S5.2f.

In conclusion, when applying the filtering strategy in (28), `kallisto|bustools`, `alevin-fry` and `STARsolo` with the `exact` and `1MMDir` strategies yielded counts from which `Seurat` was able to distinguish the biologically meaningful *red* and *PT cone PhR* clusters. However, `STARsolo` using the `1MM` strategy (designed to produce results that are near-identical to those of `Cell Ranger`), did not yield counts from which `Seurat` was able to distinguish these clusters. This reproduces the results previously reported (28), and perhaps hints at a specific mechanistic factor in the clustering differences. In particular, these results suggest that the UMI deduplication strategy plays an important role in the separation of these clusters, even when the alignment strategy itself is held fixed.

Next, we decided to investigate the effect of applying a different filtering strategy to select the cells and generate the `Seurat` count object. Specifically, the `emptyDrops` function of the `DropletUtils` package implements a procedure explicitly designed to model the ambient background distribution of expression and to select, with some userdefined false discovery rate (FDR), the barcodes corresponding to truly-captured cells. Thus, keeping the subsequent filtering, clustering and marker detection procedures the same as in the Shainer and Stemmer (28) manuscript, we filtered the barcodes using the `emptyDrops` method with default parameters, and created the resulting `Seurat` object using default parameters (Figure S5.3 and Figure S5.4). When applying this filtering, *none* of the tested methods yielded distinct clusters for the *red+* and *PT+* cells at either of the tested resolutions (0.9 and 1.2), and we also noted an attenuated signal for the epithelial cell cluster under `STARsolo` and `alevin-fry` based on the predefined markers (28). Again, while inspection of the placement of the corresponding cells in the respective UMAP embeddings, and the genes that they express, suggest that the signal separating these clusters is present in all tested methods, the clustering procedure used by `Seurat` did not separate these clusters.

Additionally, we evaluated the clustering results for the different quantification methods restricting the set of analyzed cells (i.e. barcodes) to those selected by `alevin-fry`’s knee-distance filtering procedure. Specifically, in this experiment we quantified the data with `alevin-fry`, using the knee-distance method to determine the permit-list. Subsequently, the count matrices for each method were subset to include only the barcodes appearing in this permit-list. This filtering strategy is more conservative than those examined above (i.e. fewer cells passed filtering). However, when examining the set of cells selected by this approach, all of the tested methods discovered distinct *red+* and *PT+* clusters at both the 0.9 and 1.2 resolution parameters (Figure S5.5 and Figure S5.6). This was true even for `STARsolo`’s quantification results when using the (default) `1MM` UMI resolution policy. Seemingly, this filtering of cell barcodes resulted in a signal between these clusters that was somehow clearer with respect to the method used by the `FindClusters` algorithm of `Seurat`.

Taken together, these results suggest that the main factors in the separation of these clusters during processing is a combination of the specific filtering parameters used to retain cell barcodes, in conjunction with what specific UMI deduplication strategy was used and what specific thresholds are selected for filtering. Of the methods tested, `alevin-fry`, `kallisto|bustools`, and `STARsolo` (with the `1MMDir` and `exact` UMI resolution strategies) recovered the distinct clusters under the filtering described by Shainer and Stemmer (28) and under the knee-distance filtering of `alevin-fry` (though not under the filtering that directly made use of `emptyDrops`). Conversely, `STARsolo` (with the default `1MM` UMI resolution strategy) produced counts under which these clusters were only discovered when evaluating the subset of cells returned by `alevin-fry`’s knee-distance filtering procedure. Thus, while different pre-processing methods produce counts that, under the same filtering procedures, yield distinct clustering results, there is a general tendancy for more strict filtering to elucidate a signal between these clusters that can be detected by `Seurat`’s clustering algorithm.Nonetheless, the signal itself (in terms of the biologically-relevant *opn1lw1* (31) and *parietopsin* (30) marker genes) is quite strong in the quantifications produced by all of the tested methods, if explicilty sought out. That is, by visualizing (e.g. via a `FeaturePlot`) the expression of these genes overlaid on the t-SNE embedding, there are clear subsets of the *cone PhR* cells that express one or the other of these markers, even when `Seurat` does not find them as separate clusters. This suggests that the specific clustering algorithms used may be yet another important component of the ability to automatically separate these distinct clusters of cells. Though we have not investigated the effect of this variable here, it may be an interesting direction for further work.

Finally, we return to the case of the strong DEG marker signal of the *col14a1b* gene found between the the *red* and *PT cone PhR* clusters in the `kallisto|bustools` quantifications, but which is completely absent from the filtered counts in the `STARsolo` and `alevin-fry` quantifications. To the best of our knowledge, there is no immediate or well-known biological mechanism that would cause this gene to be a differential marker between *red+* and *PT+* cells. Thus, we decided to perform a detailed read-level analysis on the expression of this gene to explore the causes of this quantification difference between `kallisto|bustools` and other tools. While the analysis described below is computationally-intensive, and therefore not feasible at scale across experiments or as a standard part of pre-processing pipelines, it helped elucidate the mechanism at work and to understand why such differences might manifest.

Specifically, we ran `kallisto` (3) in bulk mode to produce a pseudobam file that reports which underlying reads were being pseudoaligned to the constituent transcripts of *col14a1b*. When we extracted these reads and attempted to align them to the corresponding transcripts, we discovered that they almost universally produced poor quality alignments, where the only long contiguous matches between the read and the transcript were stretches of low-complexity sequence close to the indexed k-mer size (31 in this case) in length. This explains why `STARsolo`, discussed above, and `Cell Ranger`, discussed in (28), were not reporting these as valid alignments to this gene.

To explore the likely origin of these reads, we ran BLAST (32) to query these reads against the NCBI nucleotide database. For the reads we examined, the top BLAST hits contained the *pde6hb* (phosphodiesterase 6H, cGMP-specific, cone, gamma, paralog b) gene, which clearly has biologically plausible expression in this dataset. However, this gene does not appear in the Ensembl 101 *D. rerio* annotation that was used in this section and in (28). Thus, in this case, both `STARsolo` and `alevin-fry` avoided misattributing the seemingly large number of reads actually arising from *pde6hb* to other genes within the annotation, while `kallisto|bustools` falsely attributed many of these reads to *col14a1b*, for which there does not appear to be any evidence of expression.

While the spurious expression of genes when making use of a pseudoalignment-to-transcriptome approach has been previously reported by Kaminow et al. (9), and this spurious expression has even been reported to result in the expression of biologically implausible genes (33), it is of particular note in this dataset, since it results in the spurious expression of a gene that is detected as the strongest marker between these clusters of interest. The analysis here demonstrate that the strong DEG signal inferred between these clusters when using the `kallisto|bustools` quantifications is almost certainly a spurious result that derives from its use of pseudoalignment-to-transcriptome and a lack of filtering its mapping results. More generally, such occurences may not be particularly rare, and so caution should be applied when interpreting statistics like total gene detection, or median gene or UMI count, particularly among methods that employ pseudoalignment-to-transcriptome for fragment mapping, as larger values of such quantities may indicate reduced precision and not just increased sensitivity.

### 3.3. RNA Velocity of a pancreas dataset

The USA mode of `alevin-fry` generates unspliced (U), spliced (S) and ambiguous (A) count for each gene within each cell. In the pancreas dataset, the ratio of the total unspliced counts to the total spliced counts to the total ambiguous counts (U:S:A) is 0.806 : 0.125 : 0.069 over all genes within all cells. As usually RNA velocity estimators (19, 20) take only spliced and unspliced counts as the input, the ambiguous counts need either to be discarded or to be apportioned toward spliced and unspliced counts. We tested 7 different strategies for handling these ambiguous counts and observe that assigning the ambiguous count differently leads to distinct velocity and latent time estimation. In this section, we discuss the result of assigning all ambiguous counts as spliced counts, since this coincides with the reasonable prior belief that most reads in this type of experiment should arise from spliced transcripts. By doing so, the ratio of the total unspliced count to the total spliced count is then 0.875 : 0.125. The streamlines in the velocity graph (Figure 3) portray the cycling nature of the Ductal cells and endocrine progenitors, the cellular development process of endocrine progenitors (indicated by the concentration of the transcription factor Ngn3), and the differentiation process of endocrine cells, which ends with Beta cells at the latest time point, as described by Bergen et al. (20).

**Figure 3:**
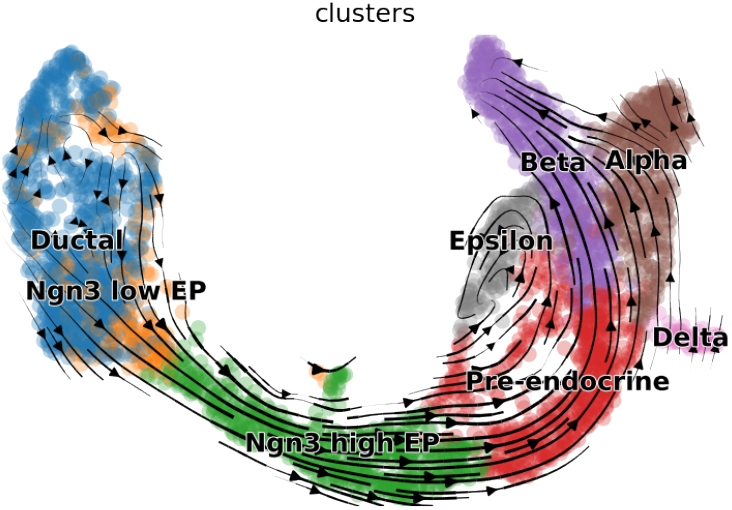
A visualization of the velocity estimation derived from `alevin-fry` counts in a UMAP-based embedding after assigning all ambiguous counts as spliced counts; the streamlines represent the velocity estimated by `scVelo`. Points (cells) are colored according to the cell type annotation.

Setting the corresponding RNA velocity related flags, for `STARsolo`, `--soloFeatures Gene Velocyto`, and for `kallisto|bustools`, `--workflow lamanno`, return the counts required by the RNA velocity pipelines. The resulting U:S:A ratio of the `STARsolo` counts is 0.834 : 0.122 : 0.044, and the resulting U:S ratio of `kallisto|bustools` counts is 0.819 : 0.181 (`kallisto|bustools` does not report ambiguous counts). Although the ratios are similar across the results of all methods, the velocity and the latent time estimation are distinct. In the velocity graph produced by `kallisto|bustools` counts (Figure S6.5a), for example, the streamlines form a back-flow, and the arrows point from the differentiated cells (Epsilon, Beta, and Alpha cells) back towards the pre-endocrine cell cluster, corresponding to the results reported in Soneson et al. (14). The velocity graph derived from `STARsolo` counts after assigning ambiguous counts to spliced (Figure S6.3b) avoids such back-flow, but does not seem to reveal the cycling population of Ductal cells, and some streamlines over the Beta cell cluster point to the opposite direction, against other streamlines over the same cell population.

Additionally, while the latent time assignments computed by `scVelo` when using the `alevin-fry` (Figure S6.2b) and `STARsolo` (Figure S6.4c) counts match the streamlines in their respective velocity graph and the latent time assignment obtained from the pre-computed counts provided in the `scVelo` tutorial, the latent time assignment derived from the `kallisto|bustools` counts is discordant with those of the other methods as well as with the directions of velocity arrows leading from the Ductal cell cluster and pre-endocrine cell cluster to the differentiated cells. Specifically, when utilizing `kallisto|bustools` counts, the latent time estimated by `scVelo` (Figure S6.5b) originates in the cluster of Beta cells, and concords with the velocity arrows leaving this cluster, but runs opposite to the main flow from the Ductal, Ngn3 and Pre-endocrine clusters into the differentiated cell clusters.

In summary, comparing the velocity graphs generated by all three methods on the endocrine pancreas dataset, the velocity streamlines and latent time assignments derived from `alevin-fry` counts well delineate the cellular development process of pancreatic endocrinogenesis, and those derived from `STARsolo` recapitulate most of the expected biology, but differ in some details, while the results derived from the `kallisto|bustools` counts only recapitulate parts of the expected biology.

### 3.4. Processing of a mouse placenta single-nucleus RNA-seq dataset

Like single-cell RNA-sequencing, single-nucleus RNA-sequencing technology is increasingly used to explore many types of biological questions, particularly in situations where full-cell scRNA-seq would be difficult or dissociation unlikely to succeed. In this section, we analyzed an snRNA-seq dataset from mouse placenta (23) that is further described in Section 2.7.

After filtering, 10,577 barcodes had (`emptyDrops`) FDR < 0.01 in the quantifications produced by `alevin-fry`. Among those, 10,483 barcodes’ mitochondrial count were lower than 0.25% and had 500 − 4, 000 expressed genes (Figure S8.1). PCA was then performed on the normalized and scaled RNA count of the top 2,000 variable genes. Based on p-values from the `JackStraw`, the first 35 PCs were selected to build the nearest neighbor graph and find cell clusters. A total of 17 clusters were found with a clustering resolution parameter of 0.6. To assign cell types for each cluster, a preprocessed `Seurat` (24) object was downloaded from the supplementary files of Marsh and Blelloch (23) (Section S8), and the cells therein were used as the reference for cell type classification using `Seurat`’s anchor transfer functionality. In this `Seurat` object, cells are classified as belonging to 5 major cell types; blood cells, decidual stroma, endothelial, fetal mesenchyme, and trophoblast. Those cell types correspond to the basic structure of the placenta, which consists of the maternal decidua, the junctional zone, and the labyrinth zone (23, 34).

By transferring the cell type annotations from the reference `Seurat` object to the `alevin-fry` result, all five clusters were detected, and the t-SNE embedding of the `alevin-fry` counts is similar to that of the reference object (Figure 4). This process was also performed for the result of `STARsolo` (Figure S8.2b) and `kallisto|bustools` (Figure S8.2c), and the five essential cell types were also detected. In conclusion, all three methods were able to retain the most significant biological signals captured in the single-nucleus RNA-seq experiment, and subsequently produced similar cell type assignment and t-SNE embedding.

**Figure 4:**
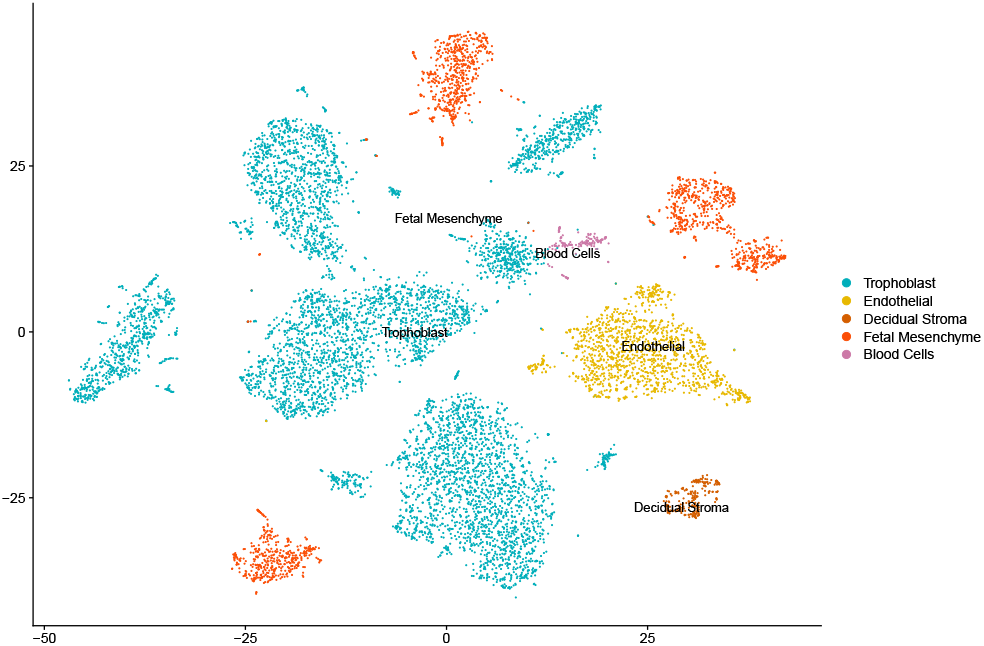
The t-SNE embedding plot of an `alevin-fry` processed mouse placenta single-nucleus dataset. Samples were filtered to remove empty droplets. RNA counts were normalized and scaled. PCA was performed on the top 2, 000 variable genes. Clustering was performed on the significant PCs. The color of each nucleus represents the inferred cell type annotation, which was learned from a reference dataset.

Subsequently, the 7,027 nuclei assigned as trophoblast in `alevin-fry` result were then selected to analyze refined trophoblast subclusters. Again, a `JackStraw` p-value informed the use of the top 28 PCs to find subclusters in the trophoblast nuclei. As some cell types have only a few corresponding nuclei, we set the clustering resolution very high (as 2.5) to detect the detailed clustering assignments and 27 clusters were found. After referring to anchors from the reference result (23), which defined 13 cell types, 12 of them were assigned to those 27 clusters. After applying the same procedure, the 6,837 trophoblast nuclei resulted in the discovery of 11 cell types in `kallisto|bustools`’s result and the 6,631 trophoblast nuclei in the `STARsolo` count matrix resulted in 10 cell types being found. Of the 13 reference clusters, `alevin-fry` counts had no resulting annotation for *SynTII precursor*, `kallisto|bustools` counts had no resulting annotation for *SynTII precursor* or *SynTI precursor*, and `STARsolo` had no resulting annotation for *SynTI precursor, LaTP* or *JZP1*. The reference labels that were not assigned across methods generally had low barcode counts in the reference dataset. It is important to note here that, just as with the cluster analysis explored in Section 3.2, the “absence” of a cluster depends on the details of the filtering approach, intermediate processing, and clustering parameters, and so the lack of a distinct cluster annotated via reference transfer does not necessarily indicate that the relevant biological signal was not present in the counts produced by a method.

In summary, all three tools demonsrated robust recapitulation of the major expected biological signals from this single-nucleus experiment, with `alevin-fry` recovering slightly more known cell types when subclustering trophoblast nuclei. Further, as outlined in Section 3.5, `alevin-fry` processed this data much faster and required considerably less memory than either `STARsolo` or `kallisto|bustools`.

### 3.5. Speed and memory usage

Finally, we assessed the speed and memory requirements of the three tools tested in this manuscript across the datasets explored in the previous sections as well as using the PBMC10k dataset (35) with the latest 10x reference annotation. We exclude `Cell Ranger` from this analysis, as it has previously been demonstrated that `STARsolo` can produce results that are almost identical to those of `Cell Ranger`, but that it is much faster and requires less RAM (9).

All experiments were conducted on a server with an Intel Xeon CPU (E5-2699 v4) with 44 cores and clocked at 2.20 GHz, 512 GB of memory, and 8 (non-RAID) 3.6 TB Toshiba MG03ACA4 HDDs, and samples were processed using a Nextflow (36) workflow. All tools tested here provide multithreaded capabilites and were run with 16 threads. All experiments were run using `STARsolo` version 2.7.9a, `salmon` v1.5.1, `alevin-fry` v0.4.0, `kb_python` 0.26.0 (with `kallisto` 0.46.2 and `bustools` 0.40.0). `STARsolo` was run with the `--soloFeatures Gene` option to process single-cell samples, with the `--soloFeatures GeneFull` option to process single-nucleus samples, and with the `--soloFeatures Gene Velocyto` option to process samples for RNA velocity analysis. `STARsolo` offers the ability to use either a dense or sampled suffix array index. Here, all tests were run using the dense suffix array, which provides the fastest runtime but which also requires more memory. If memory is at a greater premium, users can instead choose to use the sparse index, which reduces the index size by a factor of ∼ 2 but which required ∼ 1.7 times as long for processing, on average (9). `Kallisto|bustools` was run using the `kb_python` wrapper with `--workflow standard` used for single-cell samples, `--workflow nucleus` used for single-nucleus samples and `--workflow lamanno` used to process samples for RNA velocity analysis. `alevin-fry` was run in USA-mode on all samples using the `--cr-like` UMI resolution method and the appropriate counts were extracted from the resulting matrix depending upon the sample type. `alevin-fry` was tested with both the sparse and dense index as well as using both the unfiltered permit-list and filtering of barcodes prior to quantification using the knee-distance method.

Among the methods tested, `alevin-fry`, when using `sketch` mode, is always the fastest (Figure 5). When processing single-cell data and indexing only the spliced transcriptome, `kallisto|bustools` is the second-fastest tool. When both `alevin-fry` and `kallisto|bustools` are configured to use the spliced transcriptome alone as the mapping target, `alevin-fry` exhibits the lowest memory usage, followed by `kallisto|bustools`. The speed of `STARsolo` matches that of `kallisto|bustools` when the number of threads grows (often at around 16 to 20 threads depending on the specific details of the hardware configuration being used), but, by virtue of aligning against the entire genome, it consumes more memory when performing a standard (spliced) single-cell analysis. As expected, when `alevin-fry` is configured to use the *splici* transcriptome rather than just the spliced transcriptome, there is a moderate increase in th memory usage (e.g. to ∼ 10GB in dense mode and ∼ 6.5GB in sparse mode for the most recent 10X Genomics annotation of the human transcriptome). The runtime sees little effect when mapping against the *splici* transcriptome compared to the spliced transcriptome, and there also appears to be little consistent difference in the mapping speed of `alevin-fry` when using the sparse rather than the dense index. Thus, while mapping against the *splici* transcriptome requires more memory, it has little effect on the runtime and yields markedly more accurate counts, as it avoids the pitfalls of pseudoalignment-to-transcriptome described by Kaminow et al. (9).

**Figure 5:**
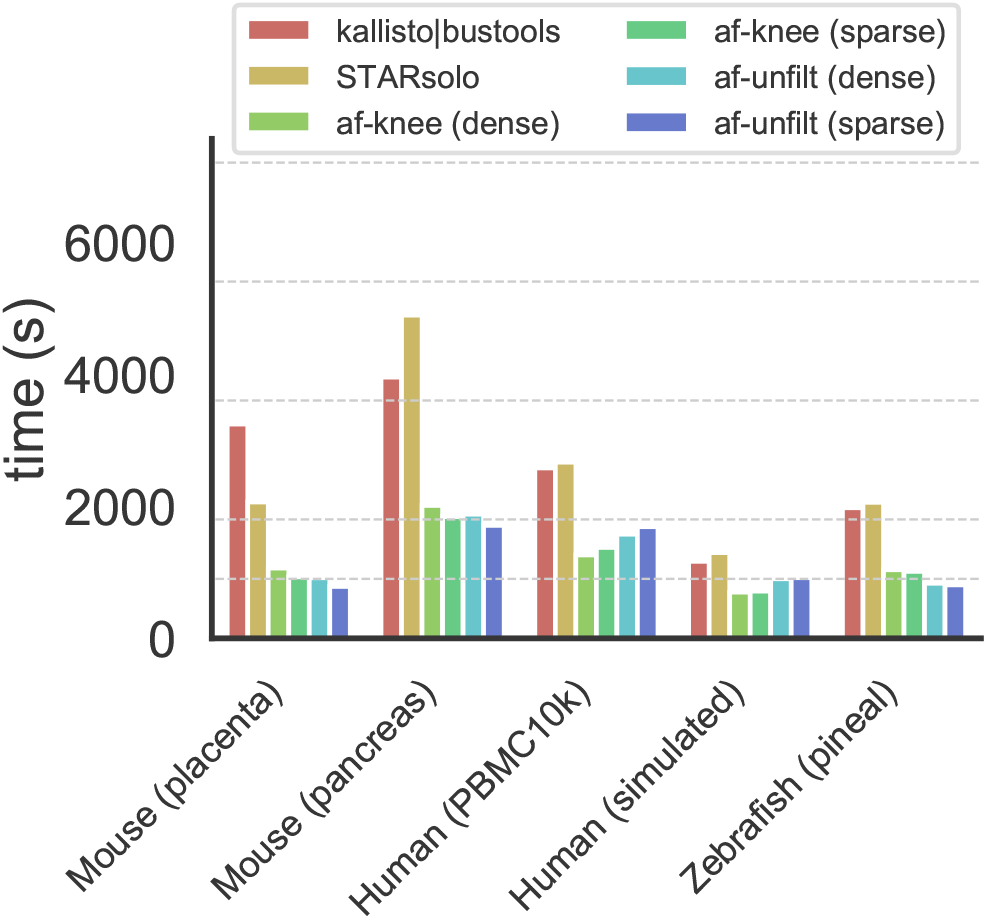
Timing for all tools (run with 16 threads) on the different datasets evaluated in this manuscript. Dashed horizontal lines appear to denote 15 minutes, 30 minutes, 60 minutes, 90 minutes and 120 minutes.

When processing single-nucleus data, `alevin-fry` is the fastest and most memory-frugal method (Figures 5 and 6). Since `STARsolo` and `alevin-fry` indices already contain the relevant intronic sequence, their index sizes do not grow when processing single-nucleus samples or preparing RNA velocity results. However, when processing single-nucleus data, there is a notable performance inversion between `STARsolo` and `kallisto|bustools`. The size of the `kallisto|bustools` index grows much larger than those of the other tools, and the speed decreases substantially. Thus, depending upon the specific organism and annotation complexity, when processing single-nucleus samples, `STARsolo` is the second-fastest and second most memory-frugal tool (even when using its dense suffix array index). On the dataset examined here, compared to `alevin-fry` (sparse, unfiltered), `STARsolo` takes ∼ 2.6 times as long and uses ∼ 6.3 times as much memory while `kallisto|bustools` takes ∼ 4.1 times as long and uses ∼ 13.1 times as much memory.

**Figure 6:**
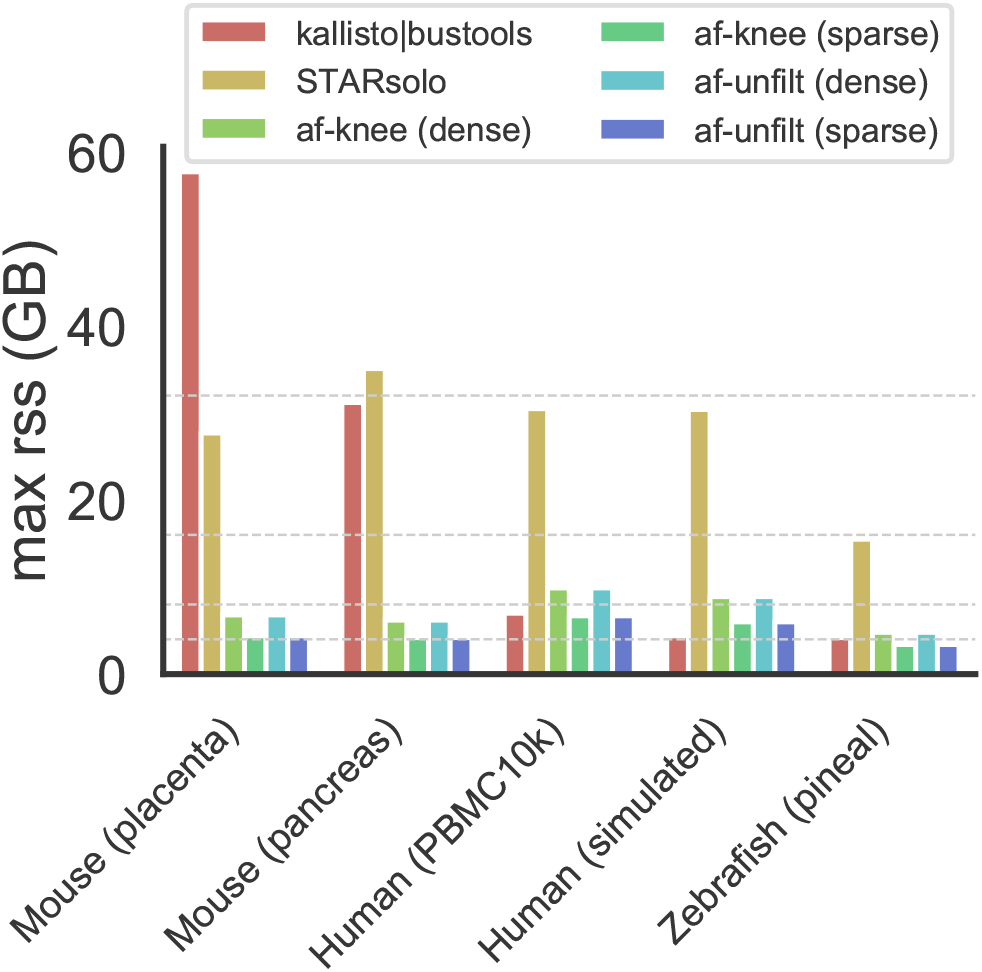
Peak memory usage for all tools on the different datasets evaluated in this manuscript. Dashed horizontal lines appear to denote 4GB, 8GB, 16GB and 32GB.

In summary `alevin-fry` is the fastest method, on average completing in under half the time required by the next fastest method. It also exhibits tightly-controlled peak memory requirements, with processing using the sparse index completing in less than 8GB of memory for all the different organisms and datasets processed in this paper. Among `STARsolo` and `kallisto|bustools`, which method is faster or which requires less memory depends on the specific type of data being processed and the details of the reference being used.

## 4. Conclusions

We have introduced `alevin-fry` as an accurate, computationally-efficient, and lightweight framework for the processing of both single-cell and single-nucleus RNA-seq data, and we have compared `alevin-fry` to both `STARsolo` and `kallisto|bustools`. `Alevin-fry` is consistently the fastest of these tools and is able to process datasets, on average, in less than half the time taken by the other tools. At the same time, when taking advantage of its sparse index, `alevin-fry` is able to process both single-cell and single-nucleus data using less than 8GB of RAM. The *splici* index, that we propose to use for all types of quantifications covered here, allows the application of a fast mapping method (pseudoalignment (3) with structual constraints) while avoiding the estimation of spurious gene expression that is observed when such approaches are applied only to the spliced transcriptome (9). This allows `alevin-fry` to quantify expression with considerably increased precision compared to other lightweight tools like `kallisto|bustools`, while using appreciably less memory than `STARsolo`.

Moreover, coupling the *splici* index with a UMI resolution method that is aware of the splicing status of different indexed targets, we introduce USA mode quantification which unifies single-cell, single-nucleus and RNA velocity preprocessing with `alevin-fry`. At the same time, `alevin-fry` is highly-configurable, providing flexibility to users at many stages of the preprocessing pipeline. For example, at the expense of a higher runtime (though not increased peak memory usage), even more precise quantifications can be obtained by performing selective-alignment (15) of sequenced fragments. Similarly, multiple options are provided for barcode (i.e. cell) permit listing and UMI resolution. `Alevin-fry` can also be used for processing other types of experiments, such as spatial scRNA-seq data and feature barcoded scRNA-seq data, and we are maintaining a growing suite of tutorials at https://combine-lab.github.io/alevin-fry-tutorials/.

We believe that `alevin-fry` strikes a remarkable balance between the often competing criteria of accuracy, performance, and flexibility, and that these characteristics make it a very appealing choice for preprocessing the rapidly-growing collection of high-throughput single-cell and single-nucleus RNA-seq data.

## 5. Data availability

The mouse pancreatic endocrinogenesis scRNA-seq dataset analyzed during the current study are available in the Gene Expression Omnibus (GEO) repository, under accession number GSM3852755 (https://www.ncbi.nlm.nih.gov/geo/query/acc.cgi?acc=GSM3852755). The mouse placenta snRNA-seq dataset analyzed during the current study are available in the GEO repository, under accession number GSM4609872 (https://www.ncbi.nlm.nih.gov/geo/query/acc.cgi?acc=GSM4609872). The *Danio rerio* pineal dataset analyzed during the current study are available in the GEO repository, under accession number GSM3511193 (https://www.ncbi.nlm.nih.gov/geo/query/acc.cgi?acc=GSM3511193).

The simulated data can be generated using the simulation scripts present in https://github.com/dobinlab/STARsoloManuscript with the PBMC5K dataset available at https://support.10xgenomics.com/single-cell-gene-expression/datasets/3.0.2/5k_pbmc_v3. The human PBMC10k dataset analyzed during the current study are available at 10X Genomics website (https://support.10xgenomics.com/single-cell-gene-expression/datasets/3.0.0/pbmc_10k_v3).

## 6. Code availability

`Alevin-fry` is written in Rust (https://www.rust-lang.org/), and is available under the BSD 3-Clause license, as a free and open source tool at https://github.com/COMBINE-lab/alevin-fry. The generation of RAD files is implemented as part of the `alevin` command of the `salmon` tool, available at https://github.com/COMBINE-lab/salmon.

Both tools are also available through bioconda (37). Useful scripts and functions for preparing a *splici* reference for indexing and for reading `alevin-fry` output in Python and R is available at https://github.com/COMBINE-lab/usefulaf. The scripts used to perform the analyses in this manuscript are available at https://github.com/COMBINE-lab/alevin-fry-paper-scripts/.

