## Supplementary Material for "Alevin-fry unlocks rapid, accurate, and memory-frugal quantification of single-cell RNA-seq data"

### S1. Some details about RAD files

The RAD files consumed by `alevin-fry` serve an essentially equivalent purpose to the BUS (1) files used by `kallisto|bustools` and are also similar to the intermediate representation of cellular barcodes and mapped UMIs used by `STARsolo` and the `dumpBfh` mode of `alevin`. Specifically, making use of this intermediate file separates the process of mapping the fragments to the target sequences from the subsequent phases of UMI resolution and quantification. Given that mapping the fragments to the target sequence is typically the computational bottleneck in preprocessing this data, the benefits of separating these tasks is manifold, but a few of the primary benefits are that (i) it allows processing the data with different filtering and resolution strategies without having to re-map the fragments (ii) it allows streaming processing of the initial FASTQ files, so that the files need not be examined more than once (so that e.g. they could be read directly from the network) and (iii) it enables low-memory processing of the data since the index can be removed from memory before subsequent processing (as is done in `kallisto|bustools` and `STARsolo`) and since the mapped records can then be re-arranged to process cells independently (as is done in `kallisto|bustools`).

However, despite serving a similar or equivalent purpose to these other files, the RAD format has some variations in design from both of these other formats. Unlike the intermediate format used by `STARsolo` and `alevin`, RAD files are not intended *just* as temporary intermediate files. Rather, they are written to disk and intended as a persistent representation of the mapped reads, potentially for multiple types of downstream processing. Unlike the BUS (1) format which encodes fixed-length records by factorizing transcript equivalence classes into a second file, the RAD format instead adopts variable length records and a generalized tagging system, similar to the SAM (2) format.

This design increases the flexibility of RAD files at the expense of potentially reducing their storage density and increasing the complexity of parsing. To recover highly-efficient and parallelizable parsing under a format that uses variable length records, RAD files adopt a chunked file format, where each chunk has a header that designates the next segment of the file that can be safely copied into memory while ending on a record boundary. Such a design has been shown to scale very well in a highly-parallel environment (3) as it separates file reading from parsing.

Additionally, unlike the BUS format, the RAD files do not contain a "factorized" representation of transcript equivalence classes. Rather, each record lists the transcripts (and bit-packed orientation) in the target index to which the corresponding fragment mapped. As mentioned above, this can result in a larger overall file size, as repeated mapping patterns are recorded independently when they occur. However, this representation also offers some benefits. First, it precludes having to maintain a synchronized global set of equivalence classes during the mapping phase, which means this structure never needs to be held in memory during mapping and any overhead associated with synchronizing it can also be avoided. Second, this decision makes it easy to separate orientation information from the transcript content of equivalence class labels. This is useful so that, during subsequent phases of processing, the mapped records can easily be subset (e.g. in the collated RAD file) to those that match the prescribed orientation, without the need to maintain or update any type of factorization structure.

### S2. Procedure for constructing a *splici* reference sequence

The term *splici* is shorthand for *spliced* + *intronic* reference sequence. This reference sequence is prepared by extracting the spliced transcripts from the reference genome according to the desired annotation (e.g. unfiltered, or filtered for certain classes of transcripts), as well as the collapsed intervals corresponding to the introns of genes. The intronic sequences of the *splici* reference play important roles in the various kinds of experiments discussed in this paper. For scRNA-seq data, although one typically focuses on the fragments and UMIs arising from the (spliced) transcriptome,and only considers the spliced and ambiguous counts when performing downstream analyses, the intronic sequences act similarly to decoy sequences proposed by Srivastava et al. (4). They account for fragments that might otherwise selectively-align or pseudoalign to the transcriptome with lower quality, but that, in fact, derive from some unspliced RNA transcript molecule. In Section 3.2, we showed that *splici* improves the accuracy and false discovery of mapping, as the intronic seuqences of *splici* act as the decoy sequence to absorb reads map to the spliced transcriptome with low quality. For snRNA-seq and RNA velocity analysis, the usage is straighforward, as the intronic sequences enable `alevin-fry` to detect the signals from unspliced, immature RNA transcripts, which are crucial in those analyses.

To construct the *splici* reference, one provides a genome fasta file and a gene annotation GTF file of the target species, as well as the read length of the experiments to be processed. Here we describe the procedure to extract the splici sequence of one gene; the procedure will be applied for all genes specified in the provided GTF file. First, the spliced transcript sequences are extracted for all isoforms of the gene. Next, the intronic regions of each isoform of the gene are extracted. These intronic regions consist of the full length of the unspliced transcript, subtracting out any exonic sequence. If alternative splicing positions occur within exons, certain sub-intervals of a gene can appear as part of both spliced transcripts and as intronic sequence. Additionally, each extracted intronic interval is extended by the provided flanking length at its starting and ending position. By adding this flanking length, the reference can account for reads that map to the junction of an exon and an intron. Otherwise these reads cannot be confidently mapped to either the unflanked intronic region or to the spliced isoforms of the gene.

The intronic regions of different isoforms often overlap considerably. To avoid repetitive indexing of shared sub-intervals, the flanked intronic regions across all isoforms of a gene will be combined if they share overlapping sub-regions. As the result, each unique genomic sequence will appear only once in the combined intronic regions of the gene. The genomic sequences of those combined intronic regions will then be extracted as the intronic part of this gene. These combined intronic sequences will each be given a unique name, and all will be added to the *splici* reference.

The process described above will then be applied to all genes defined in the provided GTF file. These sequences will be combined with any custom (user-provided) sequences, for example, one can include the sequence of mitochondrial genes to avoid spurious mapping further. The resulting sequences constitute the *splici* reference. Note that sometimes one genomic region may belong to more than one gene. In this case, extracted sequences can appear multiple times in *splici* reference with different names. As duplicated sequences will anyway appear only once in the `salmon` index except if one specifically sets the `--keepDuplicates` flag (which we have not), we did not explicitly deduplicate repeated sequences when constructing the *splici* reference. However, our script to construct the *splici* reference accepts an optional argument, `dedup_seqs` that will perform this identical sequence deduplication during reference construction.

Currently, the *splici* index depends on both the reference genome and annotation to be indexed, as well as the length of the reads that will be mapped against this index. The read length is used to determine an appropriate flanking size (default of read length $-5$) to add to the ends of the extracted collapsed intron sequences. The read length used for construction need not exactly match those being mapped, and we have evaluated that the index is reasonably robust to similar read lengths and degrades gracefully as the indexed and mapped read lengths diverge. Further, the short-read sequencing by synthesis most commonly employed for single-cell and single-nucleus experiments comprises a small set of characteristic read lengths. Nonetheless, by simply constructing the index with a single flanking length that is *as large as* the longest reads to be considered, and by propagating the relative splice junction annotations to the mapping step, it should be possible for the index to be constructed independently of the read length parameter. We are currently exploring the optimal implementation of this enhancement.

An R script that can be used to construct the *splici* reference sequences is provided in the usefulaf github repository (`https://github.com/COMBINE-lab/usefulaf`). This script uses the `eisaR` (5), `Biostrings` (6), `BSgenome` (7) and `GenomicRanges` (8) R/Bioconductor packages.

### S3. The effect of flank-length in the *splici* reference on scRNA-seq quantification

To examine the sensitivity of `alevin-fry`'s quantification results to the specific flank-length chosen when building the *splici* reference sequence, we built a *splici* sequence for the reference genome used in Section 3.1, using flank lengths from 85 nucelotides up through 145 nucleotides (in increments of 5 nucleotides). We then quantified the simulated fragments and evaluated the mean cell-level Spearman correlation, the MARD (dropping NA values), and the mean relative false-positive and false-negative expression rate per cell. We find that the quantification results are robust to the specific flank length chosen Table S3.1. We observe a slight improvement in the accuracy as the flank-length increases which might be due to detecting more reads mapping to the intronic regions, we note that this trend may only be valid for the standard single cell rna-seq data and probably not in the context of RNA-velocity analysis. While it is important to consider these empirical results in context — they are obtained from simulated data and in the context of scRNA-seq quantification and not e.g. RNA-velocity analysis — they nonetheless provide evidence for the robustness of the quantification results to the selected flank length.

| **flank-length** | mean Sp. corr. | MARD (drop NA) | mean rFP/cell | mean rFN/cell |
|---|---|---|---|---|
| **85** | 0.9883 | 0.0263 | 0.0110 | 0.0118 |
| **90** | 0.9883 | 0.0263 | 0.0109 | 0.0118 |
| **95** | 0.9883 | 0.0263 | 0.0109 | 0.0118 |
| **100** | 0.9884 | 0.0262 | 0.0109 | 0.0118 |
| **105** | 0.9884 | 0.0262 | 0.0109 | 0.0118 |
| **110** | 0.9884 | 0.0262 | 0.0109 | 0.0118 |
| **115** | 0.9884 | 0.0262 | 0.0108 | 0.0118 |
| **120** | 0.9884 | 0.0262 | 0.0108 | 0.0118 |
| **125** | 0.9884 | 0.0261 | 0.0108 | 0.0118 |
| **130** | 0.9884 | 0.0261 | 0.0107 | 0.0118 |
| **135** | 0.9885 | 0.0260 | 0.0107 | 0.0118 |
| **140** | 0.9885 | 0.0260 | 0.0107 | 0.0118 |
| **145** | 0.9885 | 0.0260 | 0.0107 | 0.0118 |

Table S3.1: The quantification performance on the simulated human data set as a function of the flank-length used to construct the *splici* reference sequence.

### S4. Alternative UMI resolution strategies

Currently, the `cr-like` and `cr-like-em` methods, described in Section 2, are exposed in both the USA-mode of `alevin-fry` tested in this manuscript, as well as the splice-unaware quantification mode that does not differentiate among targets based on their splicing status. Additionally, `alevin-fry` offers two other UMI resolution methods that work for splice-unaware quantification but that are not currently enabled in USA-mode. These other methods may provide useful capabilities for UMI resolution, and that we anticipate enabling these for USA-mode quantification in future releases of `alevin-fry`. We describe these alternative UMI resolution methods below.

**Parsimony resolution** The parsimony based resolution, enabled with the `parsimony` argument to the resolution option of `alevin-fry`, implements part of the UMI resolution strategy originally proposed by Srivastava et al. (9). The algorithm starts by collapsing the input record information for each cell into a set of tuples that encode transcript-level equivalence class labels, UMIs and associated counts. That is, each tuple consists of a set of transcript identifiers to which the corresponding fragment mapped, the UMI associated with these fragments, and the number of times this transcript set / UMI pair was observed.

Next, a parsimonious UMI graph (PUG) is constructed, in which each node consists of one the tuples described above (i.e. (equivalence class label, UMI, count)). Nodes are connected by a bidirected edge if their equivalence class labels have a non-null intersection and they share an identical UMI. Additionally, nodes are connected by a directed edge if their equivalence class labels have a non-null intersection, their UMIs have an edit distance of 1, and the count associated with one endpoint is at least twice the count associated with the other endpoint. The edge is directed from the node with the higher count to the node with the lower count. Following the intuition of Smith et al. (10), the PUG structure connects in the graph those UMIs that are likely candidates to have arisen from a single pre-PCR mRNA molecule, and where differences in UMI sequence may be the result of sequencing or amplification error.

Given the PUG constructed in this manner, the algorithm then greedily searches for a minimum cover of this PUG by monochromatic arborescences (9). Each monochromatic arborescence is a vertex disjoint subgraph of the PUG that is consistently labeled by a set of transcripts, which are the pre-PCR molecules capable of explaining the observed reads and UMIs, and their sequence relationship. Intuitively, the search for a minimum cover attempts to maximize the parsimony of the explaination for the data represented by the PUG. For each monochromatic arborescence, the algorithm checks the set of transcripts; if all transcripts belong to a single gene, all deduplicated UMIs within this arboresence will be assigned to this gene. If more than one gene is involved in this maximal monochromatic arboresence, the corresponding fragments (and UMIs) are discarded.

**Parsimony + EM resolution** If the `parsimony-em` (previously `full`) argument is provided to the resolution option of `alevin-fry`, the algorithm executed is precisely the same as what is described above, except that maximal monochromatic arboresences labeled by more than one gene are not discarded. Rather, the genes that equally explain this arboresence become the label of a gene-level equivalence class. After all records for a cell are processed, these gene-level equivalence classes and their corresponding counts are processed by an expectation-maximization algorithm, and the multi-gene reads are probabilistically allocated among the genes as in (9).

The `alevin-fry` framework makes it easy to add new UMI resolution strategies, and to test their effect on quantification independent of other steps of preprocessing. In addition to the `parsimony` and `parsimony-em` methods, we anticipate that `alevin-fry` will be a productive testing ground for alternative and improved UMI resolution methods in the future.

### S5. Additional plots for analysis of a *Danio rerio* pineal experiment.

In Section 3.2, we talked about the effects of the UMI deduplication strategy and the cell filtering strategy to the expression of marker genes. In this section, we demonstrate the expression status of the marker genes of all cell types in a dot plot manner. We also show the resulting t-SNE embedding for all the strategies.

To be specific, we tested 3 different cell filtering strategies, which are `emptyDrops` (11), the strategy proposed in Shainer and Stemmer (12), and the cells identified by `alevin-fry` knee-finding strategy. For UMI deduplication strategies, which is specific to `STARsolo` since it suffers from losing the ability to detect the expression of the marker genes the most, we tested `1MM`, `1MMDir` and `Exact`. The filtered count matrices were processed and analyzed with `Seurat` using the same pipeline described in the Section 2.7. The cell type of each cluster was determined by the marker genes described by Shainer and Stemmer (12). The expression of those marker genes under different settings and their corresponding t-SNE embeddings are shown in Figure S5.1 and Figure S5.2 for the results filtered by the Shainer and Stemmer (12) strategy, in Figure S5.3 and Figure S5.4 for the results filtered by `emptyDrops`, and in Figure S5.5 and Figure S5.6 for the results using cells identified by `alevin-fry` knee-finding strategy.

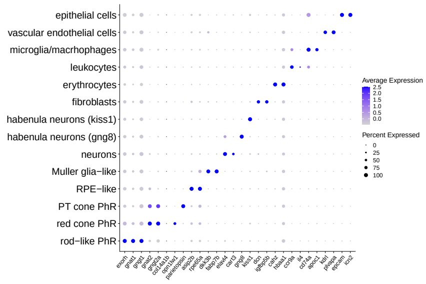


(a) `alevin-fry`, clustering resolution=0.9

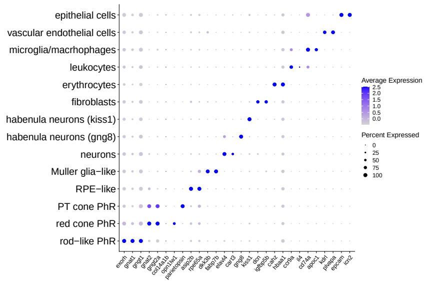


(b) `alevin-fry`, clustering resolution=1.2

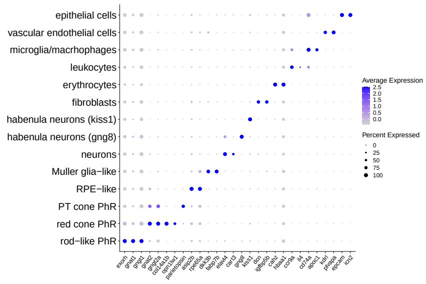


(c) `kallisto|bustools`, clustering resolution=0.9

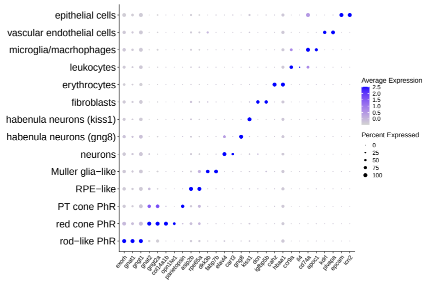


(d) `kallisto|bustools`, clustering resolution=1.2

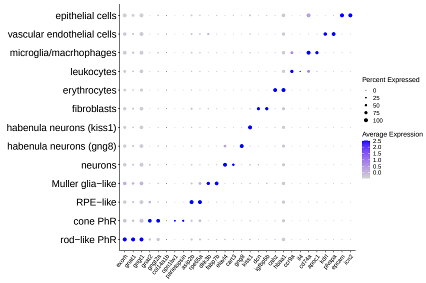


(e) `STARsolo 1mm` clustering resolution=0.9

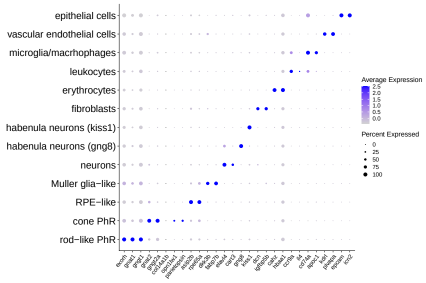


(f) `STARsolo 1mm`, clustering resolution=1.2

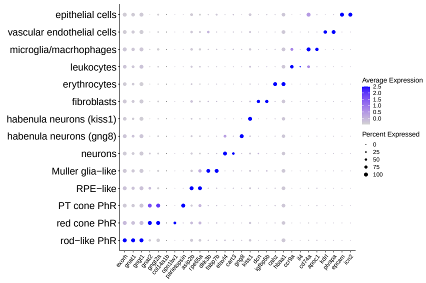


(g) `STARsolo 1mmDir`, clustering resolution=0.9

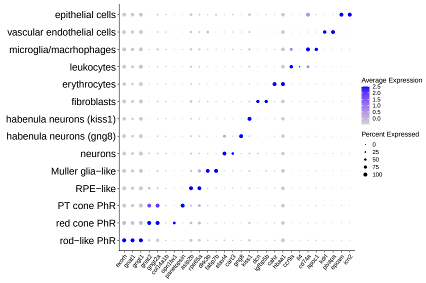


(h) `STARsolo 1mmDir`, clustering resolution=1.2

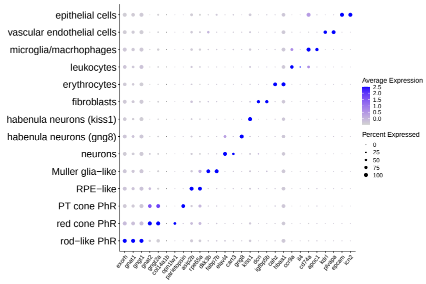


(i) `STARsolo exact`, clustering resolution=0.9

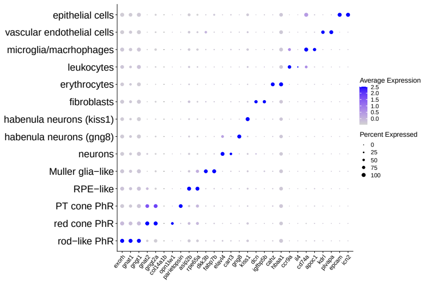


(j) `STARsolo exact`, clustering resolution=1.2

Figure S5.1: This collection shows the dot plots of the expression of the marker genes of different cell type clusters in the `alevin-fry`, `kallisto|bustools` and `STARsolo` processed *Danio rerio* pineal dataset. Samples (cells) are filtered using the strategy introduced in (12). The color of the dots represent the expression level of each marker gene within each cell type. The size of the dots represent the proportion of cells in which the marker gene is expressed. The figures in the first column used clustering resolution as 0.9. The figures at the second column used clustering resolution as 1.2.

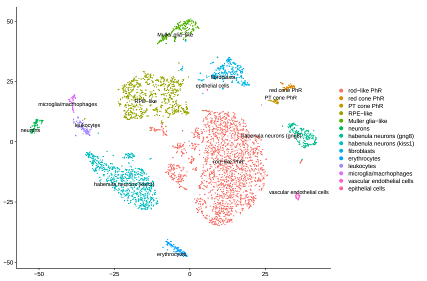

(a) `alevin-fry`, clustering resolution=0.9

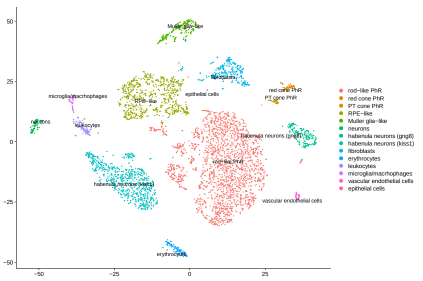

(b) `alevin-fry`, clustering resolution=1.2

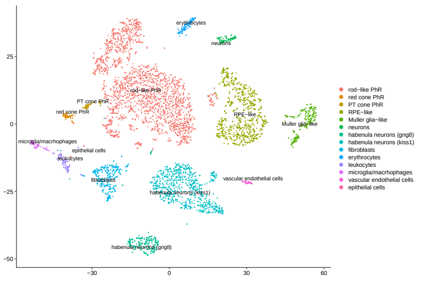

(c) `kallisto|bustools`, clustering resolution=0.9

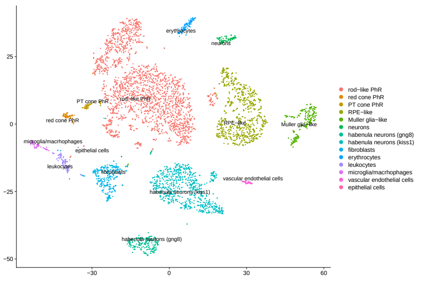

(d) `kallisto|bustools`, clustering resolution=1.2

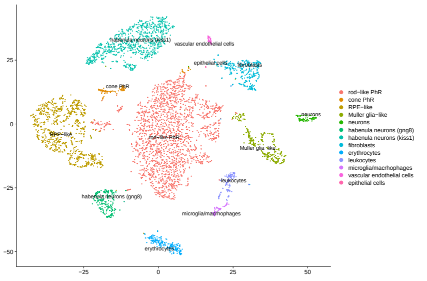

(e) `STARsolo 1mm`, clustering resolution=0.9

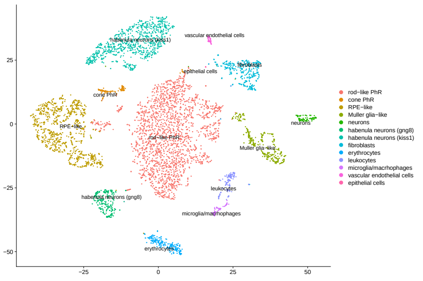

(f) `STARsolo 1mm`, clustering resolution=1.2

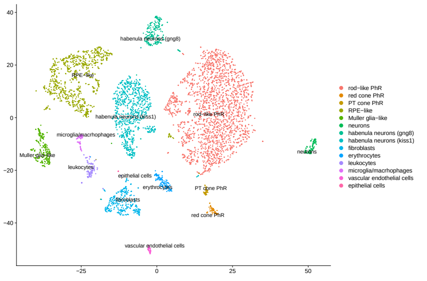

(g) `STARsolo 1mmDir`, clustering resolution=0.9

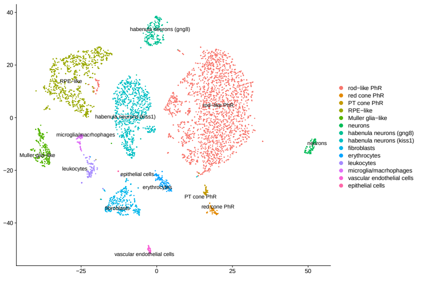

(h) `STARsolo 1mmDir`, clustering resolution=1.2

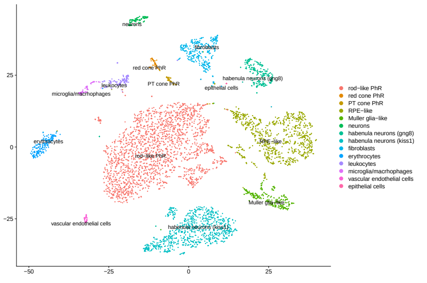

(i) `STARsolo exact`, clustering resolution=0.9

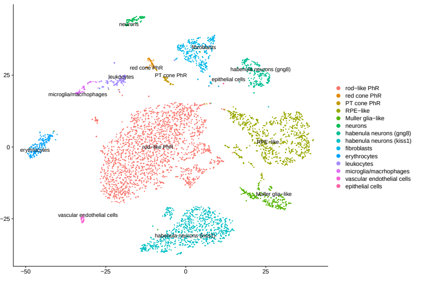

(j) `STARsolo exact`, clustering resolution=1.2

Figure S5.2: This collection shows the t-SNE embedding of the `alevin-fry`, `kallisto|bustools` and `STARsolo` processed *Danio rerio* pineal dataset. Samples (cells) are filtered using the strategy introduced by (12). Each point represents a sample (cell) in the dataset.The color of the points represent the cell type annotation of cells. The figures in the first column used clustering resolution as 0.9. The figures in the second column used clustering resolution as 1.2.

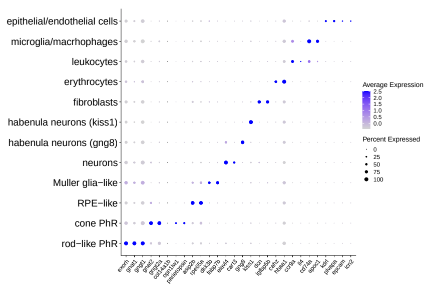


(a) `alevin-fry`, clustering resolution=0.9

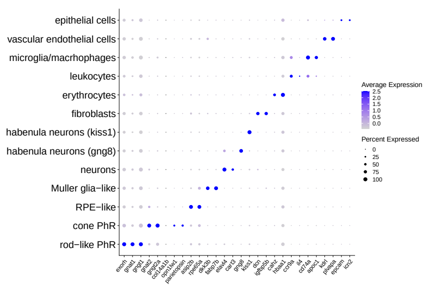


(b) `alevin-fry`, clustering resolution=1.2

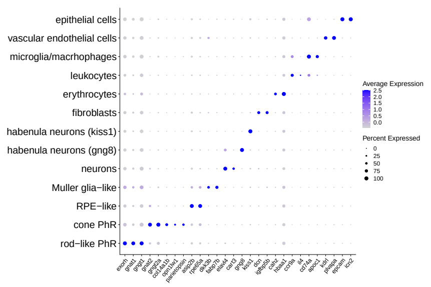


(c) `kallisto|bustools`, clustering resolution=0.9

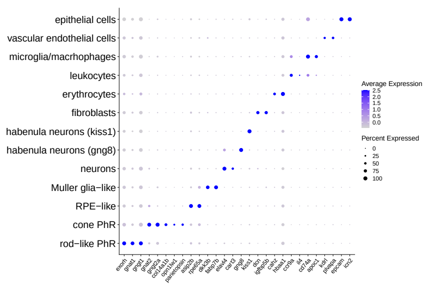


(d) `kallisto|bustools`, clustering resolution=1.2

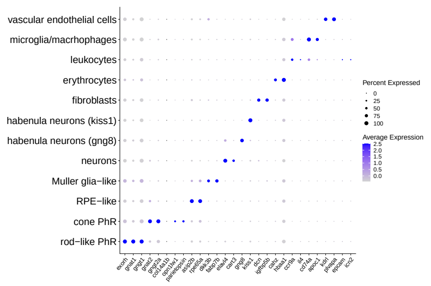


(e) `STARsolo 1mm` clustering resolution=0.9

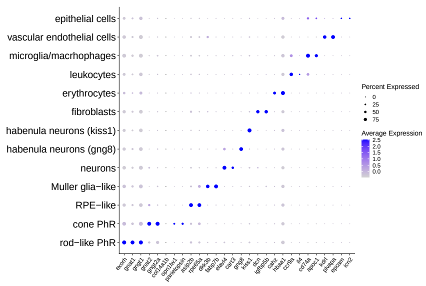


(f) `STARsolo 1mm`, clustering resolution=1.2

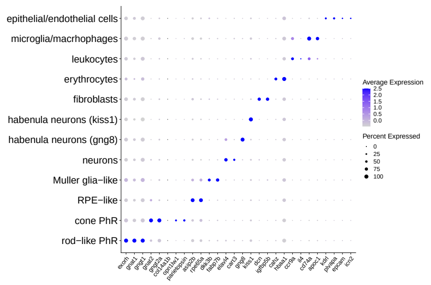


(g) `STARsolo 1mmDir`, clustering resolution=0.9

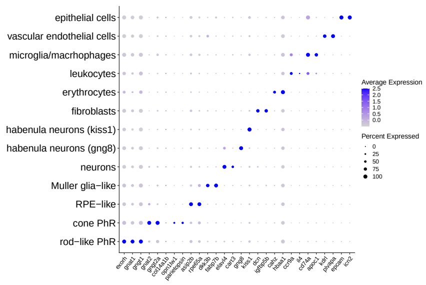


(h) `STARsolo 1mmDir`, clustering resolution=1.2

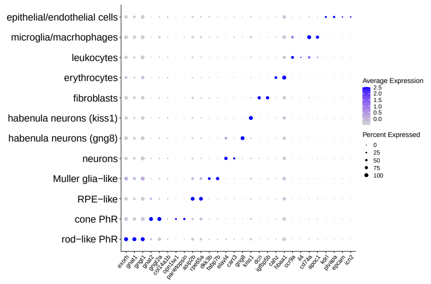


(i) `STARsolo exact`, clustering resolution=0.9

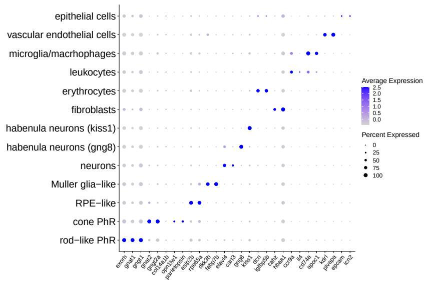


(j) `STARsolo exact`, clustering resolution=1.2

Figure S5.3: This collection shows the dot plots of the marker gene expression of different cell type clusters in the `alevin-fry`, `kallisto|bustools` and `STARsolo` processed *Danio rerio* pineal dataset. Samples (cells) are filtered using `emptyDrops` as described in the Methods section. The color of the dots represents the expression level of each marker gene within each cell type. The size of the dot represents the proportion of cells in which the marker gene is expressed. The figures in the first column used clustering resolution as 0.9. The figures in the second column used clustering resolution as 1.2.

### S5 ADDITIONAL PLOTS FOR ANALYSIS OF A *DANIO RERIO* PINEAL EXPERIMENT.

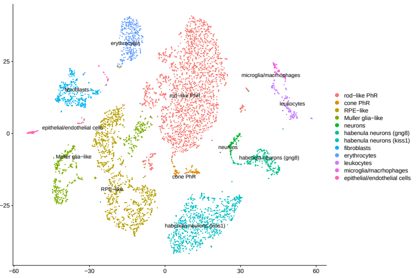


(a) `alevin-fry`, clustering resolution=0.9

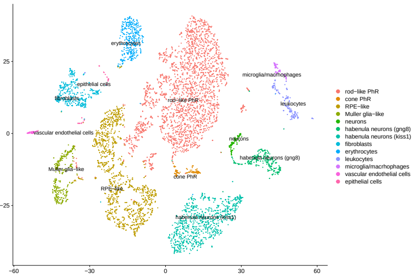


(b) `alevin-fry`, clustering resolution=1.2

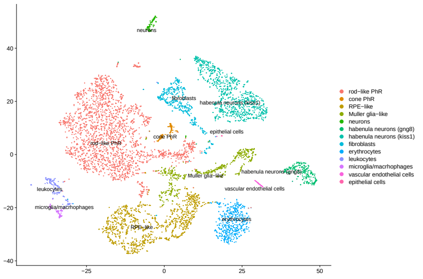


(c) `kallisto|bustools`, clustering resolution=0.9

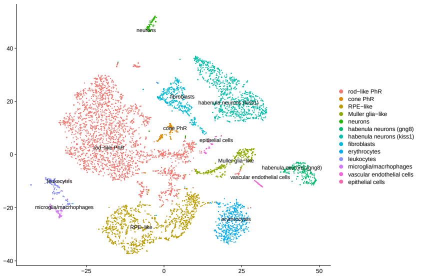


(d) `kallisto|bustools`, clustering resolution=1.2

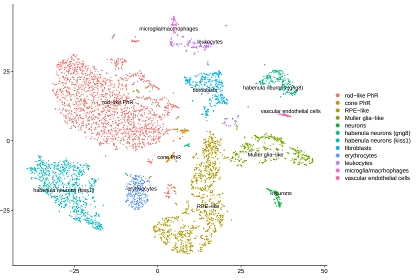


(e) `STARsolo 1mm`, clustering resolution=0.9

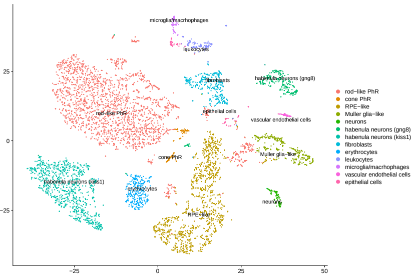


(f) `STARsolo 1mm`, clustering resolution=1.2

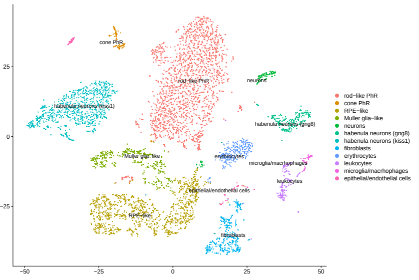


(g) `STARsolo 1mmDir`, clustering resolution=0.9

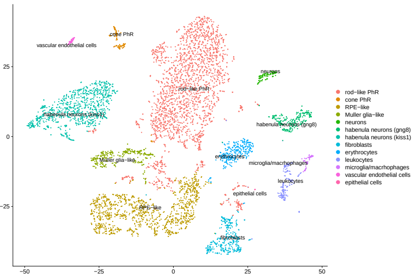


(h) `STARsolo 1mmDir`, clustering resolution=1.2

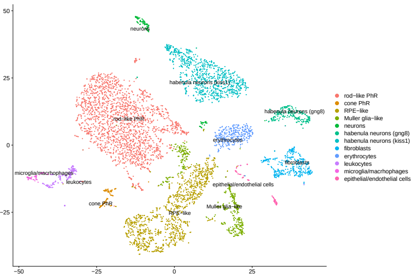


(i) `STARsolo exact`, clustering resolution=0.9

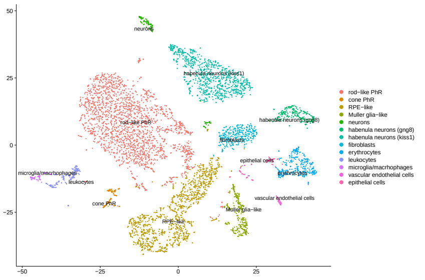


(j) `STARsolo exact`, clustering resolution=1.2

Figure S5.4: This collection shows the t-SNE embedding of the `alevin-fry`, `kallisto|bustools` and `STARsolo` processed *Danio rerio* pineal dataset. Samples (cells) are filtered using `emptyDrops` as described in the Methods section. Each point represents a sample (cell) in the dataset. The color of the points represent the cell type annotation of cells. The figures in the first column used clustering resolution as 0.9. The figures in the second column used clustering resolution as 1.2.

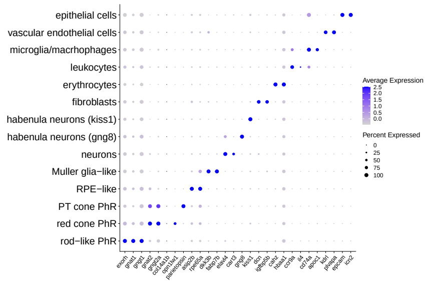


(a) `alevin-fry`, clustering resolution=0.9

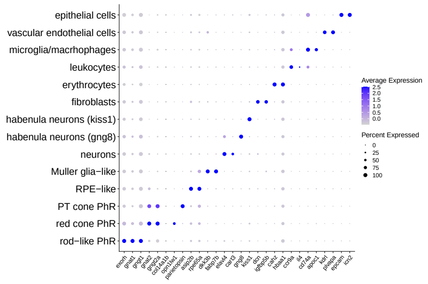


(b) `alevin-fry`, clustering resolution=1.2

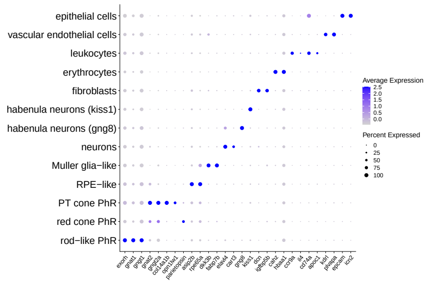


(c) `kallisto|bustools`, clustering resolution=0.9

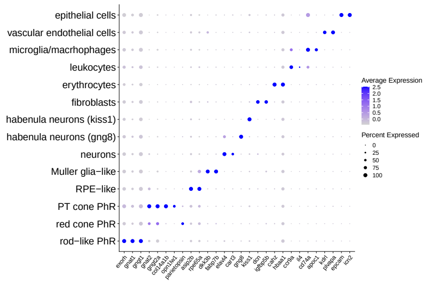


(d) `kallisto|bustools`, clustering resolution=1.2

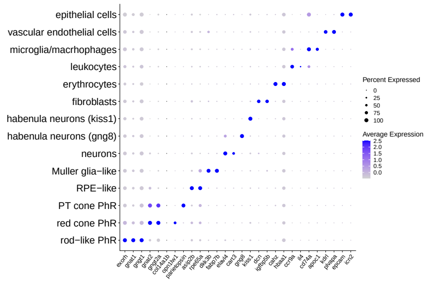


(e) `STARsolo 1mm` clustering resolution=0.9

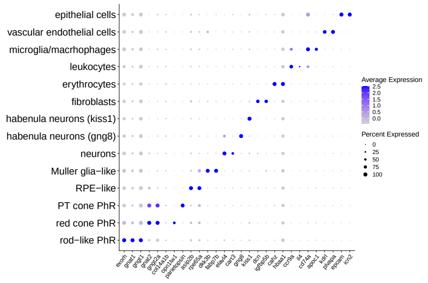


(f) `STARsolo 1mm`, clustering resolution=1.2

Figure S5.5: This collection shows the dot plots of the marker gene expression of different cell type clusters in the `alevin-fry`, `kallisto|bustools` and `STARsolo` processed *Danio rerio* pineal dataset. Samples (cells) are filtered using the cell barcode list returned by `alevin-fry` knee-finding. The color of the dots represent the expression level of each marker gene within each cell type. The size of the dots represent the proportion of cells in which the marker gene is expressed. The figures in the first column used clustering resolution as 0.9. The figures in the second column used clustering resolution as 1.2.

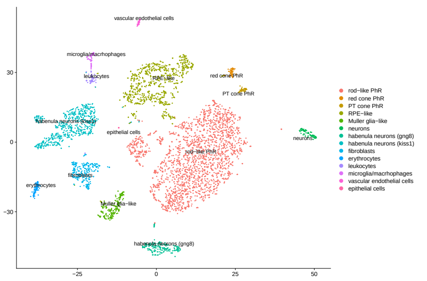

(a) `alevin-fry`, clustering resolution=0.9

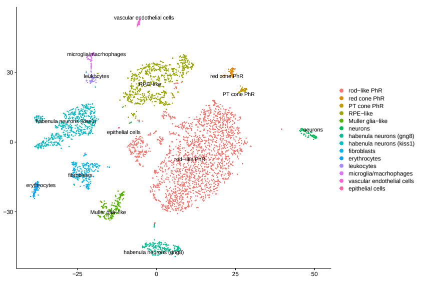

(b) `alevin-fry`, clustering resolution=1.2

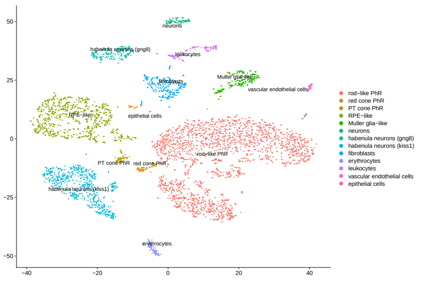

(c) `kallisto|bustools`, clustering resolution=0.9

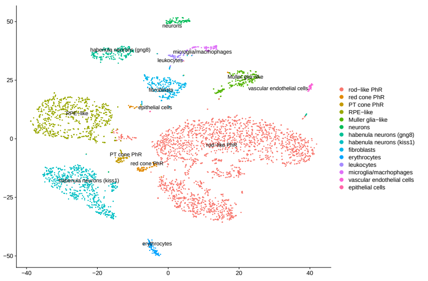

(d) `kallisto|bustools`, clustering resolution=1.2

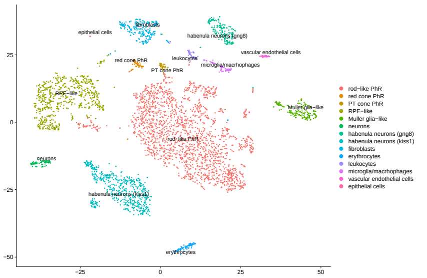

(e) `STARsolo 1mm`, clustering resolution=0.9

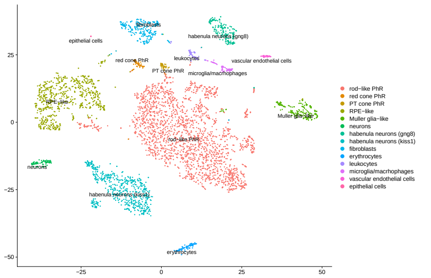

(f) `STARsolo 1mm`, clustering resolution=1.2

Figure S5.6: This collection shows the t-SNE embedding of the `alevin-fry`, `kallisto|bustools` and `STARsolo` processed *Danio rerio* pineal dataset. Samples (cells) are filtered using the cell barcode list returned by `alevin-fry` knee-finding. Each point represents a sample (cell) in the dataset. The color of the points represent the cell type annotation of cells. The figures in the first column used clustering resolution as 0.9. The figures in the second column used clustering resolution as 1.2.

### S6. Alternative ambiguous allocation strategies for RNA velocity

In this section, we provide all additional RNA velocity related figures, including the velocity streamlines of different strategies to process the ambiguous counts for `STARsolo` and `alevin-fry`, and the corresponding latent time plots. As `kallisto|bustools` returns only the spliced counts and the unspliced counts, the methods of assigning ambiguous counts affect only `STARsolo` and `alevin-fry`.

Setting RNA velocity related flags, `alevin-fry` and `STARsolo` return a spliced count, an unspliced count and an ambiguous count for each gene in each individual cell. However, `kallisto|bustools` resolves the ambiguity internally and returns only a spliced count and an unspliced count for each gene with in each cell. To run an RNA velocity analysis, which usually takes the spliced and unspliced count for each gene as input, we need to process the ambiguous counts for `STARsolo` and `alevin-fry` results.

We tested 7 different strategies to process the ambiguous counts, which are for each gene within each individual cell, (i) discarding the ambiguous counts, (ii) regarding the ambiguous count as spliced, (iii) regarding the ambiguous count as unspliced, (iv) evenly distributing the ambiguous count to spliced and unspliced, (v) dividing the ambiguous count by the ratio of confidently spliced count to the confidently unspliced count, (vi) dividing the ambiguous count by the ratio of not-unspliced (spliced + ambiguous) to unspliced, and (vii) dividing the ambiguous counts by the ratio of spliced to not-spliced (unspliced + ambiguous).

The process of running `scVelo` is as follows: The `pp.filter_and_normalize` function was applied to identify the top $2,000$ variable genes, and median normalize the count of those variable genes. The first- and second-order moments of the normalized spliced and unspliced counts of each gene were calculated via `pp.moments`. The velocity dynamics, including reaction rates and latent variables, was recovered using `tl.recover_dynamics` function. Gene specific RNA velocities were estimated using the dynamical mode by `tl.velocity`, and were visualized in a UMAP embedding using `pl.velocity_embedding_stream`. Cell specific latent time estimates were computed using `tl.latent_time` and were visualized using the `scatter` function.

The visualization of the velocity streamlines derived from the seven strategies are shown in Figure S6.1 for `alevin-fry`, Figure S6.3 for `STARsolo`, and Figure S6.5a for `kallisto|bustools`. The figures are based on the UMAP embedding provided in the `scVelo` provided example dataset. The cell type assignments are also from the `scVelo` example dataset. Each point represents a sample in the dataset. The streamlines represent the visualized RNA velocity direction. The color of points represent the cell type assignments of cells.

The estimated latent time assignment of samples are shown in Figure S6.2 for `alevin-fry`, Figure S6.4 for `STARsolo`, and Figure S6.5b for `kallisto|bustools`. The figures were built upon the UMAP embedding provided in the `scVelo` example pancreas dataset. Each point represents a cell in the dataset. points are colored according to the assigned latent time, which is a numeric value between $0$ and $1$, where $0$ represents the starting state, and $1$ represents the terminal state.

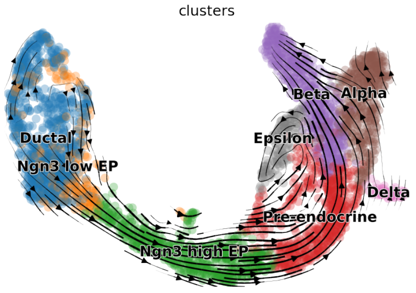


(a) Discarding ambiguous counts.

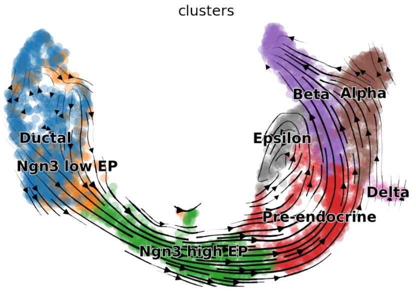


(b) Assigning ambiguous counts to spliced.

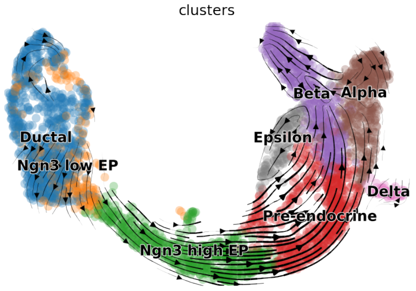


(c) Assigning ambiguous counts to unspliced.

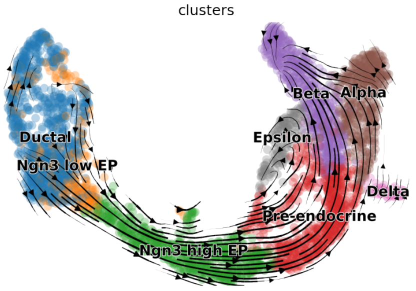


(d) Dividing ambiguous counts by the ratio of spliced to unspliced.

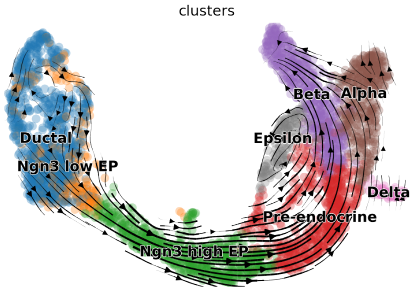


(e) Dividing ambiguous counts by the ratio of not-unspliced to unspliced.

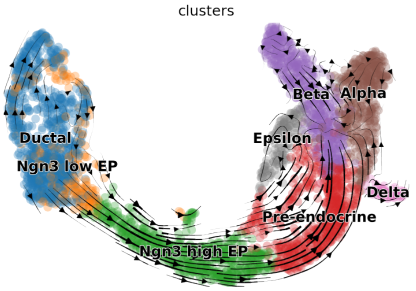


(f) Dividing ambiguous counts by the ratio of spliced to not-spliced.

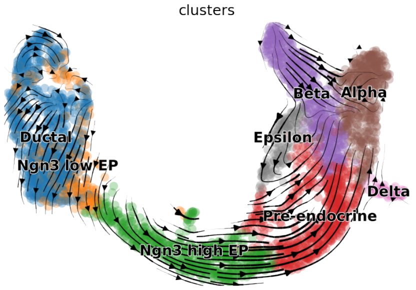


(g) Dividing ambiguous counts evenly.

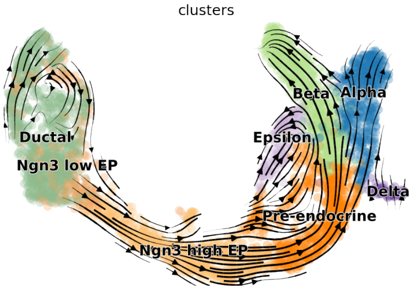


(h) scVelo provided example dataset.

Figure S6.1: The visualized velocity streamlines of the result of `alevin-fry` after assigning ambiguous counts using 7 different ways. The result returned from the `scVelo` provided example dataset is shown in (h). All figures use the same UMAP embedding derived from the `scVelo` example dataset. The arrows represent the estimated RNA velocity direction. Points (cells) are colored according to the cell type annotation.

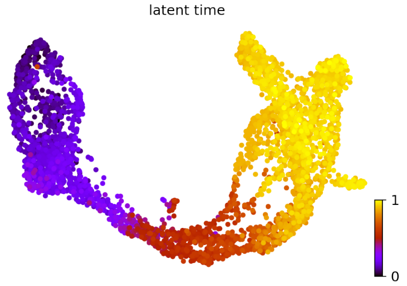


(a) Discarding ambiguous counts.

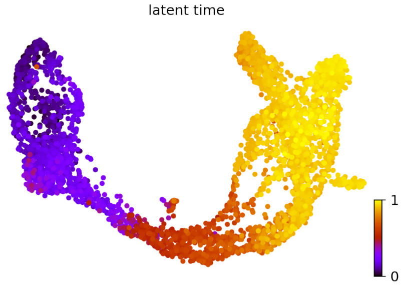


(b) Assigning ambiguous counts to spliced.

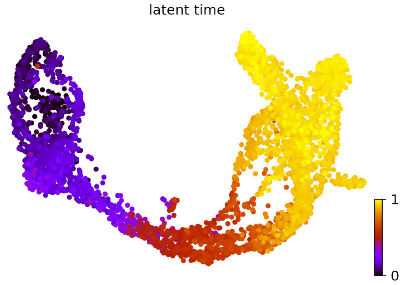


(c) Assigning ambiguous counts to unspliced.

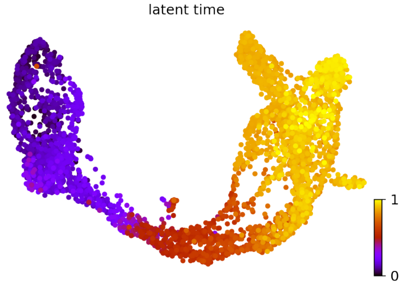


(d) Dividing ambiguous counts by the ratio of spliced to unspliced.

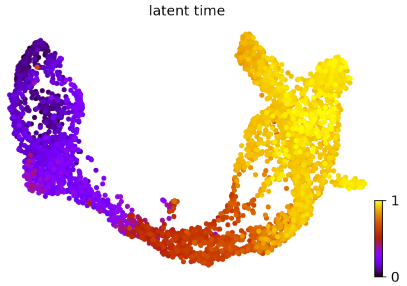


(e) Dividing ambiguous counts by the ratio of not-unspliced to unspliced.

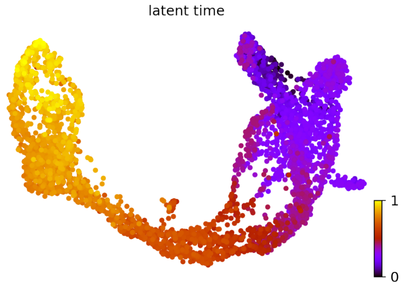


(f) Dividing ambiguous counts by the ratio of spliced to not-spliced.

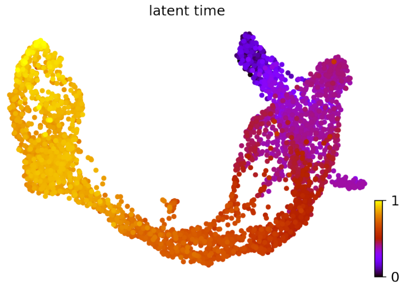


(g) Dividing ambiguous counts evenly.

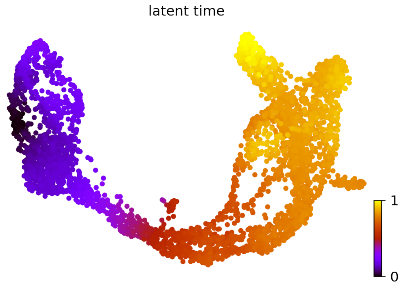


(h) scVelo provided example dataset.

Figure S6.2: The latent time assignments of the result of `alevin-fry` after assigning ambiguous counts using 7 different ways. The result returned from the `scVelo` provided example dataset is shown in (h). All figures use the same UMAP embedding derived from the `scVelo` example dataset. Points (cells) are colored according to the latent time assignments.

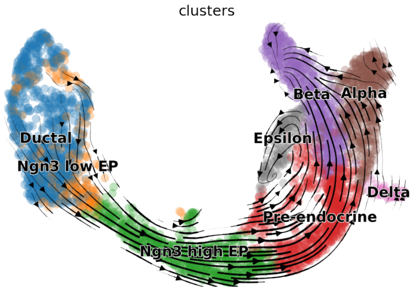


(a) Discarding ambiguous counts.

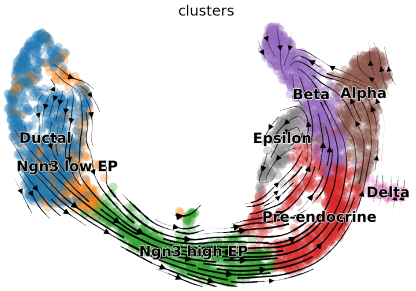


(b) Assigning ambiguous counts to spliced.

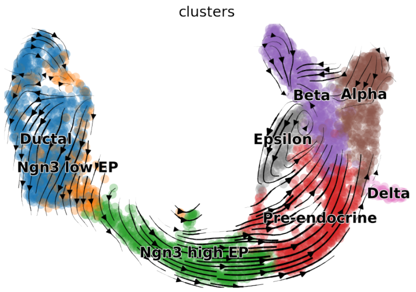


(c) Assigning ambiguous counts to unspliced.

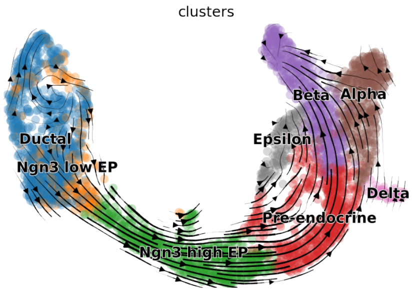


(d) Dividing ambiguous counts by the ratio of spliced to unspliced.

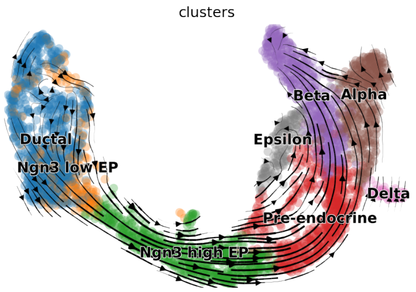


(e) Dividing ambiguous counts by the ratio of not-unspliced to unspliced.

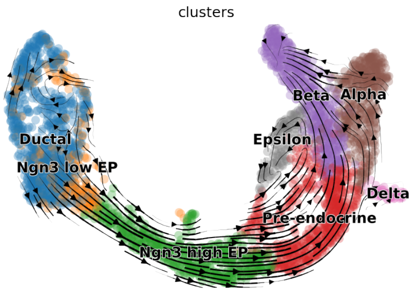


(f) Dividing ambiguous counts by the ratio of spliced to not-spliced.

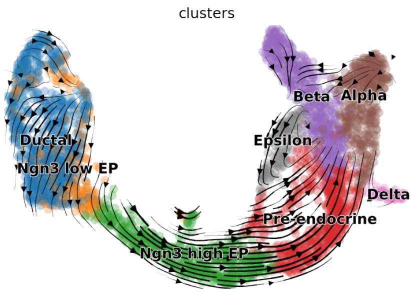


(g) Dividing ambiguous counts evenly.

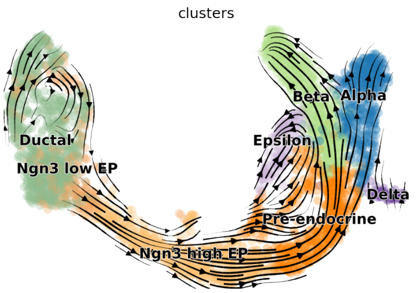


(h) scVelo provided example dataset.

Figure S6.3: The visualized velocity streamlines of the result of `STARsolo` after assigning ambiguous counts using 7 different ways. The result returned from the `scVelo` provided example dataset is shown in (h). All figures use the same UMAP embedding derived from the `scVelo` example dataset. The arrows represent the estimated RNA velocity direction. Points (cells) are colored according to the cell type annotation.

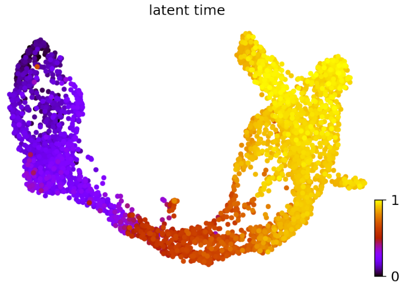


(a) Discarding ambiguous counts.

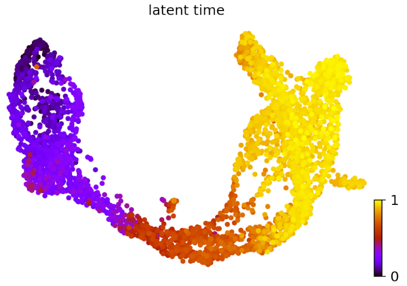


(b) Assigning ambiguous counts to spliced.

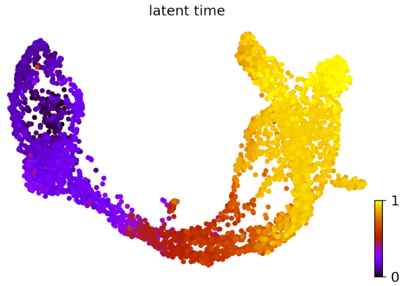


(c) Assigning ambiguous counts to unspliced.

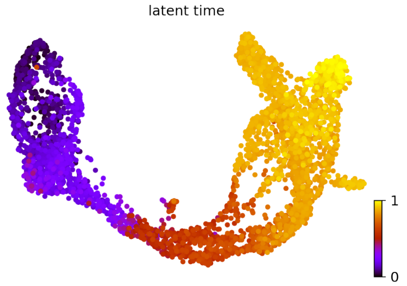


(d) Dividing ambiguous counts by the ratio of spliced to unspliced.

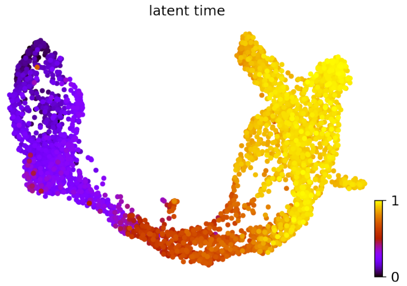


(e) Dividing ambiguous counts by the ratio of not-unspliced to unspliced.

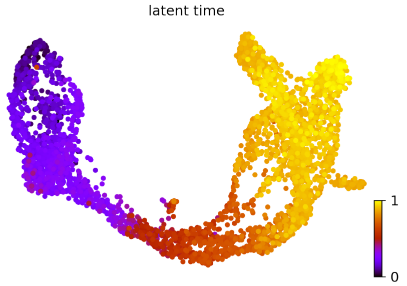


(f) Dividing ambiguous counts by the ratio of spliced to not-spliced.

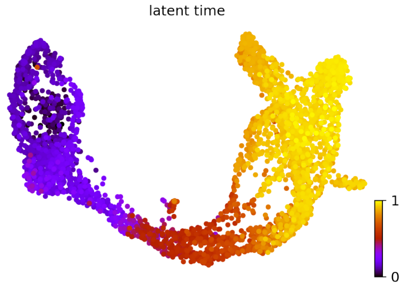


(g) Dividing ambiguous counts evenly.

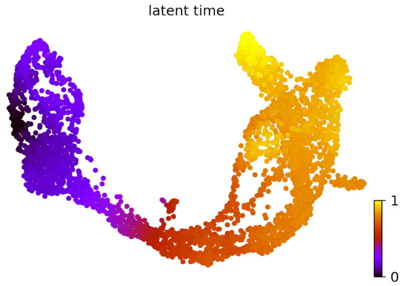


(h) scVelo provided example dataset.

Figure S6.4: The latent time assignments of the result of `STARsolo` after assigning ambiguous counts using 7 different ways. The result returned from the `scVelo` provided example dataset is shown in (h). All figures use the same UMAP embedding derived from the `scVelo` example dataset. Points (cells) are colored according to the latent time assignments.

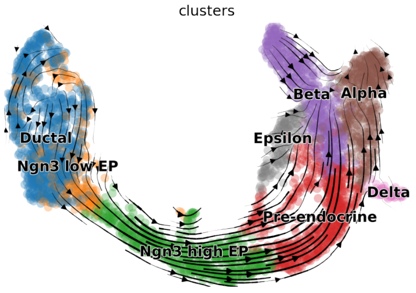


(a) The visualized velocity streamlines of the result of `kallisto|bustools` atop of the UMAP embedding derived from the `scVelo` example dataset. The arrows represent the estimated RNA velocity direction. Points (cells) are colored according to the cell type annotation.

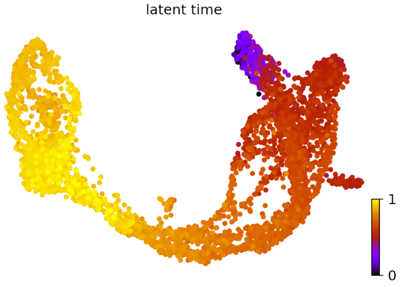


(b) The latent time assignments of the result of `kallisto|bustools` atop of the UMAP embedding derived from the `scVelo` example dataset. Points (cells) are colored according to the latent time assignments.

Figure S6.5: The visualized velocity and latent time assignment of `kallisto|bustools` result.

### S7. Simulated data evaluation metrics

In order to provide a well-rounded view of accuracy, a number of different performance metrics are reported when comparing the predictions of different tools to the ground truth simulated data. Here, we provide brief definitions of these metrics. In the equations below, $M$ is the number of genes being quantified and $C$ is the number of cells (i.e. corrected barcodes) being quantified. We compute two variants of the mean absolute relative deviation (MARD) that differ only in how division by 0 is treated. The definition of MARD used here is adopted from the manuscript of Kaminow et al. (13) and differs from how the MARD is sometimes defined by using a max operation rather than a sum in the denominator. The first variant, MARD (drop NA) is defined as follows:

$$\text{MARD}_{\text{dropNA}} = \frac{1}{E}\sum_{i=1}^{C}\sum_{j\in E_i}\frac{|x_{ij}-y_{ij}|}{\max(x_{ij},y_{ij})} \quad \textbf{(1)}$$

where $x_{ij}$ is the true count for gene $j$ in cell $i$, and $y_{ij}$ is the estimated count for gene $j$ in cell $i$.

$$E_i = \{j \mid \max(x_{ij},y_{ij})) > 0\} \quad \textbf{(2)}$$

and $E = \sum_{i=1}^{C}|E_i|$. Likewise, we can define a variant of the MARD where predictions of 0 abundance for truly unexpressed genes are counted as 0 (rather than ignored); this variant is defined as

$$\text{MARD}_{\text{NA}=0} = \frac{1}{M\cdot C}\sum_{i=1}^{C}\sum_{j=1}^{M}\begin{cases}\frac{|x_{ij}-y_{ij}|}{\max(x_{ij},y_{ij})} & \text{if } \max(x_{ij},y_{ij}) > 0\\ 0 & \text{otherwise}\end{cases}. \quad \textbf{(3)}$$

We also compute the mean relative false-positive expression per cell (rFP/cell) and the mean relative false-negative expression per cell (rFN/cell). We define

$$\text{mean rFP/cell} = \frac{1}{C}\sum_{i=1}^{C}\frac{1}{T_i}\sum_{j=1}^{M}\begin{cases}1 & \text{if } y_{ij} > 0 \text{ and } x_{ij} = 0\\ 0 & \text{otherwise}\end{cases} \quad \textbf{(4)}$$

and

$$\text{mean rFN/cell} = \frac{1}{C}\sum_{i=1}^{C}\frac{1}{T_i}\sum_{j=1}^{M}\begin{cases}1 & \text{if } y_{ij} = 0 \text{ and } x_{ij} > 0\\ 0 & \text{otherwise}\end{cases}, \quad \textbf{(5)}$$

where

$$T_i = \sum_{j=1}^{M}\begin{cases}1 & \text{if } x_{ij} > 0\\ 0 & \text{otherwise}\end{cases}.$$

It is important to note that these terms described in Equations (4) and (5) are not standard definitions of false positive and false negative, as they measure the fraction of false positives and false negatives relative to the number of *truly-expressed* genes in each cell, rather than to the total number of genes.

### S8. Clustering analysis of mouse placenta single-nucleus dataset

In this section, we provide the detailed pipeline used in the clustering analyses, and show the resulting figures derived from these analyses.

To fairly compare the results across methods, we implement the `emptyDrops CR` filtering function, which is an `R` implementation of the filtering approach implemented by `STARsolo`, which is itself a re-implementation of the method present in `Cell Ranger` (version $\geq 3$). This approach first runs a basic UMI count filter, and then defines a specific range of candidate barcodes for which `emptyDrops` is used to differentiate between valid captured barcodes and likely-empty droplets. This approach was adopted after we discovered that running `emptyDrops` without this initial filtering results in many more cells being called than are expected in the data (for all quantification methods).

After filtered empty droplets, the RNA counts of each nucleus were log normalized using `NormalizeData` from `Seurat`. Next, the top $2,000$ variable genes were detected by `FindVariableFeatures` with default parameter setting. The RNA counts of those $2,000$ variable genes were then scaled using `ScaleData` function. Then, PCA was performed with those variable genes, and the significant PCs were selected according to the p-values returned by the `JackStraw` function using a simple knee criterion based on the cumulative p-value distribution. Using those significant PCs, the t-SNE dimensionality reduction was calculated using the `RunTSNE` function, the nearest neighbor graph was constructed using `FindNeighbors` and the clusters was detected using the `FindClusters` function with clustering resolution $0.6$. In order to assign a cell type to each cluster, all samples in the R object *AllStages_AllNuclei_obj.Rdata*, which is provided as a supplementary file in Marsh and Blelloch (14) and were downloaded from https://figshare.com/projects/Single_nuclei_RNA-seq_of_mouse_placental_labyrinth_development/92354, were used as the reference to transfer the cell type annotation from the reference samples to the query object. To be specific, `FindTransferAnchors` was first applied to find the anchors between the reference and query dataset using their respective significant PCs. Then, the `TransferData` function was used to transfer the cell type annotation from the reference object to the query object according to the anchors calculated in the previous step and the significant PCs. Then, the nuclei assigned as trophoblast were then selected to explore the subclusters in the trophoblast. Similar to the previous procedure, clusters were found using the significant PCs, and cell type was learned from the R objected, *AllStages_TrophoblastNuclei_obj.Rdata*, again provided as a supplementary file in Marsh and Blelloch (14).

Next, we briefly describe the actual process. When analyzing snRNA-seq data, the unspliced, spliced and ambiguous counts for each gene in each cell returned from the USA mode of `alevin-fry` were summed to arrive at the overall per-cell count for each gene. For `kallisto|bustools` results, the spliced and unspliced count were summed to get the total count. Barcodes were filtered to remove likely-empty droplets (using the `emptyDrops CR` function), barcodes with too many or too few unique genes, and barcodes with high mitochondrial concentration. Figure S8.1 shows the number of expressed genes in each barcode (*nFeature_RNA*), the total RNA count of each barcode, and the mitochondrial concentration of each barcode of `alevin-fry` counts of unfiltered samples (Figure S8.1a) and the samples after filtering with `emptyDrops CR` and futher filtering based on the number of expressed genes and mitochondrial concentration. (Figure S8.1b). Notice that the majority of the unfiltered samples have low mitochondrial count, which is desired in scRNA-seq because mitochondria is suppose to live in the cytoplasm. After the filtering steps, the retained barcodes have reasonable expressed gene and RNA counts and low mitochondrial concentration. The remaining barcodes were regarded as properly identified nuclei and were used in the clustering analysis.

After processed by the procedure described above, all three methods resulted in similar t-SNE embedding and clustering results, and all displayed similar structure to the t-SNE embedding of the reference dataset (Figure S8.2). Notice that we also provided the t-SNE plot of the E14.5 samples in the reference dataset in panel (d). The cell type annotation of those samples were included in the reference dataset.

The nuclei assigned as trophoblast in each dataset were then selected to explore the cell types in the trophoblast. After learned the cell type assignment from the reference dataset, which defined 13 different clusters, `alevin-fry` counts resulted in 12 clusters, `kallisto|bustools` counts resulted in 11 clusters and `STARsolo` counts resulted in 10 clusters (Figure S8.3). The expression of the marker genes defined in Shainer and Stemmer (12) was shown in Figure S8.4. We also provide the t-SNE plot of the E14.5 samples in the reference dataset in Figure S8.2d. The cell type annotation of those samples were included in the reference dataset.

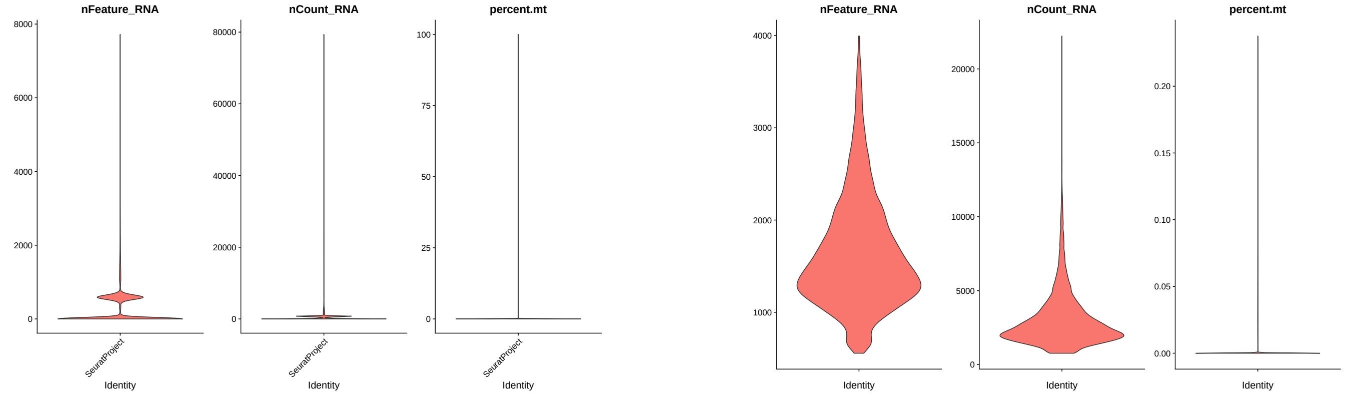


(a) QC violin plots of `alevin-fry` raw counts without any filtering.

(b) QC violin plots of `alevin-fry` result after filtering with `emptyDrops CR` and futher filtering based on the number of expressed genes and mitochondrial concentration.

Figure S8.1: Quality control plots of `alevin-fry` processed counts from the single-nucleus RNA-seq mouse placenta dataset. From left to right, the three figures show the number of expressed genes, the total RNA count and the mitochondrial concentration of samples.

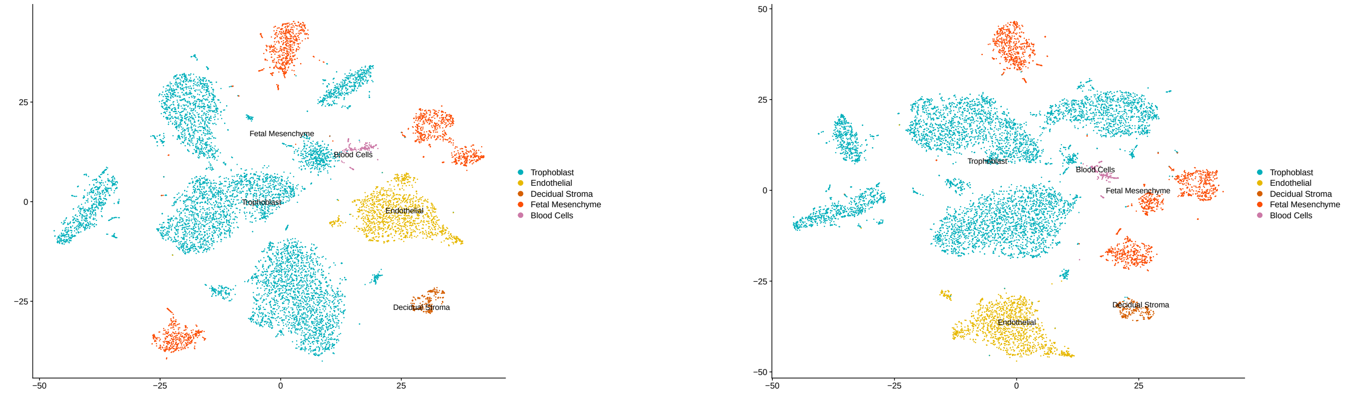


(a) `alevin-fry`

(b) `STARsolo`

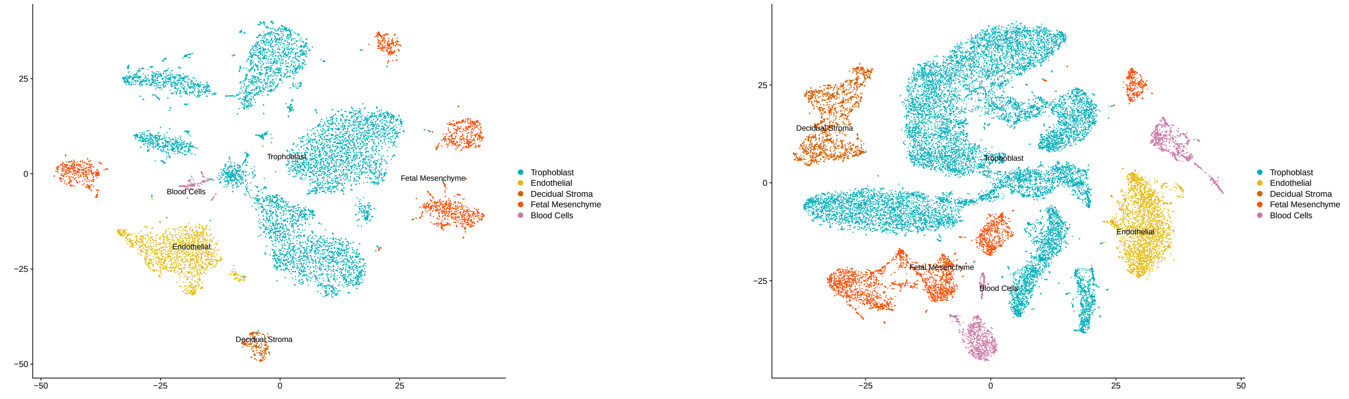


(c) `kallisto|bustools`

(d) E14.5 samples in the reference dataset

Figure S8.2: t-SNE embedding plots of `alevin-fry`, `STARsolo` and `kallisto|bustools` counts of a mouse placenta single-nucleus dataset, and the samples from E14.5 in the reference dataset. RNA counts were normalized and scaled. PCA was performed on the top 2,000 variable genes. Clustering was performed on the significant PCs. The color of each nuclei represents the inferred cell type annotation, which was learned from a reference dataset.

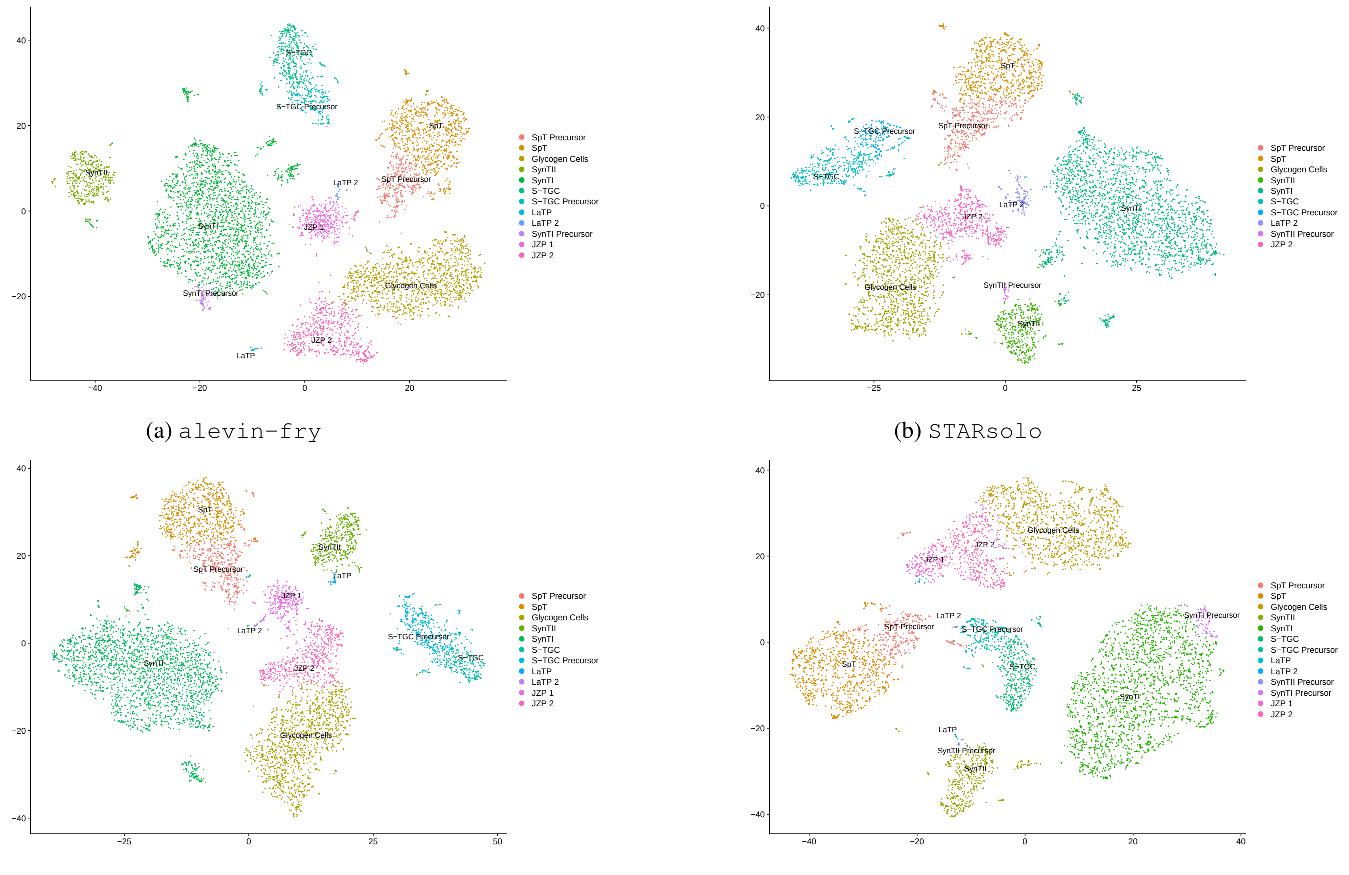


(a) `alevin-fry` (b) `STARsolo` (c) `kallisto|bustools` (d) samples from E14.5 in the reference trophoblast dataset

Figure S8.3: The clusters identified from the trophoblast nuclei in the `alevin-fry`, `STARsolo`, and `kallisto|bustools` result of a mouse placenta dataset, and the samples from E14.5 in the trophoblast reference dataset. Each point represents a nucleus in the dataset. Points are colored according to the cell type assignment.

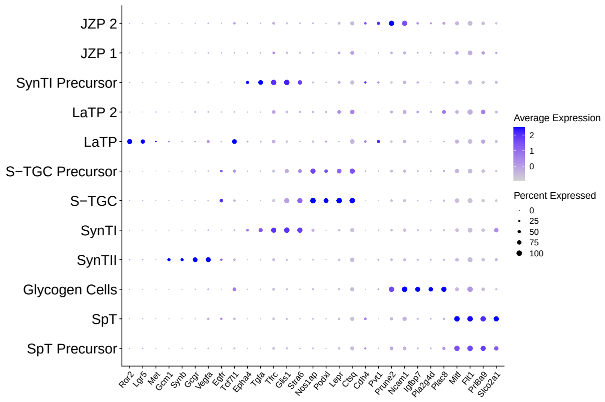


(a) `alevin-fry`

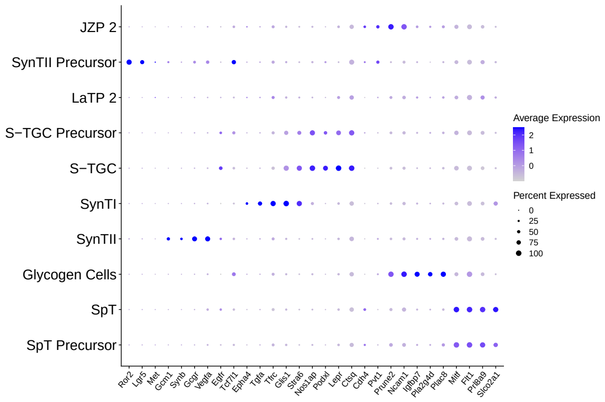


(b) `STARsolo`

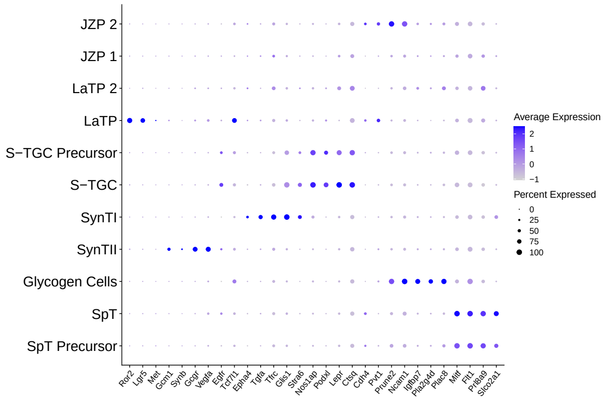


(c) `kallisto|bustools`

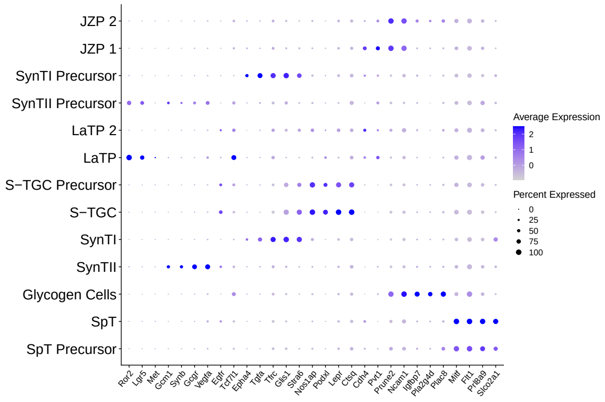


(d) The samples from E14.5 in the reference dataset

Figure S8.4: The expression level of the marker genes of each trophoblast cell type in the trophoblast nuclei of the `alevin-fry`, `STARsolo` and `kallisto|bustools` results of a mouse placenta dataset, and the samples from E14.5 in the reference object. The color of the dots represent the expression level of each marker gene within each nuclei. The size of the dots represent the proportion of cells in which the marker gene is expressed. The cell type annotations, learned from a reference dataset, appear on the y-axis.
